# SpikeLab: Agentic tools for spike data analysis

**DOI:** 10.64898/2026.04.25.720833

**Authors:** Tjitse van der Molen, Luka Cheney, Kamran Hussain, Ojas Brahme, Ash Robbins, Max Lim, Alex Spaeth, Jinghui Geng, David F. Parks, Kenneth S. Kosik, Mircea Teodorescu, David Haussler, Tal Sharf

**Author notes:** Correspondence (T.V.D.M.), (T.S.).

## Abstract

Large language models have the potential to transform scientific research and analysis, but without domain-specific structure they produce silent methodological errors, unreported decisions, and irreproducible results. Here we present SpikeLab, a text-to-analysis framework for neural spike data that combines composable data structures with a skill-based agentic system enforcing bounded autonomy: mandatory use of expert-vetted methods, correctness over efficiency, and clarification-seeking on ambiguous requests. In a controlled benchmark on electrophysiology data, Sonnet 4.6 with SpikeLab produced correct and reproducible results across all tasks, outperforming both the unassisted Sonnet and the more capable Opus 4.6, which exhibited deterministic failures including *ad hoc* method invention, silent data reduction, and inconsistent experimental designs. We demonstrate versatility across *in vivo* mouse, human, and *in vitro* brain organoid recordings, and apply the framework to a pharmacological dose-response study spanning single-unit dynamics, pairwise network structure, burst-level temporal sequences, and latent population states, all through natural language prompts without writing analysis code.

## Main

Large language models are reshaping how scientists interact with data. From autonomous chemical synthesis^1^ to mathematical discovery^2^ and multi-agent nanobody design^3^, AI systems are increasingly capable of executing complex research workflows with minimal human intervention.^4,5^ In computational biology, domain-specific foundation models and agents have been developed for single-cell multi-omics^6^ and autonomous transcriptomics analysis.^7^

Tool-augmented LLM agents have shown that pairing models with expert-designed tools outperforms unassisted models on domain tasks.^8,9,10,11^ The growing adoption of LLMs in scientific writing and analysis is now quantifiable, an estimated 22% of recent computer science papers show evidence of LLM involvement.^12^

However, the reliability of LLM-generated scientific code remains a serious concern. A benchmark of 16 LLMs on 293 biomedical data science tasks found overall accuracy below 40%.^13^ LLM-generated code suffers from systematic failure modes including fabricated API calls, incorrect or arbitrary parameter usage, and missing edge case handling.^14,15^ Critically, many of these errors are silent. For example, the code runs without raising exceptions but produces incorrect results. This is a particularly dangerous failure mode in scientific contexts where the output may be interpreted as a finding. Beyond correctness, reproducibility is also at risk. A LLM may generate syntactically valid code without specifying the dependency environments needed for reliable re-execution.^16^ Practical guidelines for maintaining scientific rigor when using AI coding assistants have begun to emerge^17^, but these address researcher behavior, rather than the tools themselves.

The flexibility that makes LLMs useful as a general tool limits their precision in science exploration. An unrestricted LLM can explore large number of possible approaches to a given problem, but this freedom allows it to invent *ad hoc* methods, make implicit assumptions, and silently simplify computations, all without reporting these choices to the user.^13,18,19,20^ The result is analysis that appears correct but may not follow established methodology, may not be reproducible across runs, and may not be comparable to results obtained through well-established, scientifically validated pipelines. A structured environment that constrains model behavior is therefore essential for ensuring correctness, and reproducibility in the analysis of biological data.

Neuroscience presents a compelling case for this approach. Extracellular electrophysiology pipelines involve a large number of interdependent analysis stages. These range from spike sorting, firing rate estimation, pairwise correlation, population dynamics modeling, each with domain-specific conventions^21^. LLMs have shown strong performance on neuroscience prediction tasks^22^ and have been applied to neurophysiology data exploration^23^, but the gap between predicting results and producing reliable analysis code remains wide. Spike sorting frameworks such as SpikeInterface^24^ and Kilosort^25^ have standardized upstream processing, yet the downstream analysis of sorted spike data, from single-unit statistics through network-level metrics to latent population dynamics, lacks equivalent standardization, leaving LLMs to improvise solutions from first principles with each invocation.

Here we present SpikeLab, a text-to-analysis framework that enables natural language-driven neural spike data analysis through combining agentic tool orchestration with expert-vetted analysis pipelines. SpikeLab provides composable data structures and analytical methods that can be assembled into analysis pipelines by researchers directly or by an agentic system responding to natural language prompts. A skill-based agentic interface enforces bounded autonomy, which include mandatory use of library methods, correctness over efficiency, and clarification-seeking on ambiguous requests. Taken together, these skill-based agentic constraints ensure that the model’s flexibility is guided by validated building blocks rather than *ad hoc* improvisation.

The combination of a robust data class foundation with a dedicated developer skill enables community contributors to integrate new analyses into the library through the same agentic workflow, expanding the framework’s capabilities over time without compromising its methodological guarantees. In this way, the SpikeLab repository functions as a persistent, expert-validated knowledge base that any analysis agent can draw from, encoding domain expertise not as transient prompt context but as reusable, tested, and documented code.

We benchmark SpikeLab against unassisted frontier models on a standardized analysis pipeline, demonstrate its versatility across *in vivo* mouse, human, and *in vitro* brain organoid recordings, and apply it to a multi-condition pharmacological case study spanning single-unit dynamics to latent population states.

## Results

### Library design

SpikeLab is a text-to-analysis framework that allows researchers to describe analyses in natural language, rather than writing analysis code directly. A skill-based agentic system translates these descriptions into executable code using the library’s structured data classes and methods. At its core, SpikeLab provides a set of stable, composable building blocks in the form of data structures and analytical methods, that can be combined to construct a wide range of analysis pipelines. These analysis pipelines can either be assembled manually by anyone calling the SpikeLab methods in Python or automatically by an agentic system responding to natural language prompts.

### Core data structures

SpikeLab is built around five core data classes that form a composable hierarchy (**Fig. 1**).

**Figure 1.**
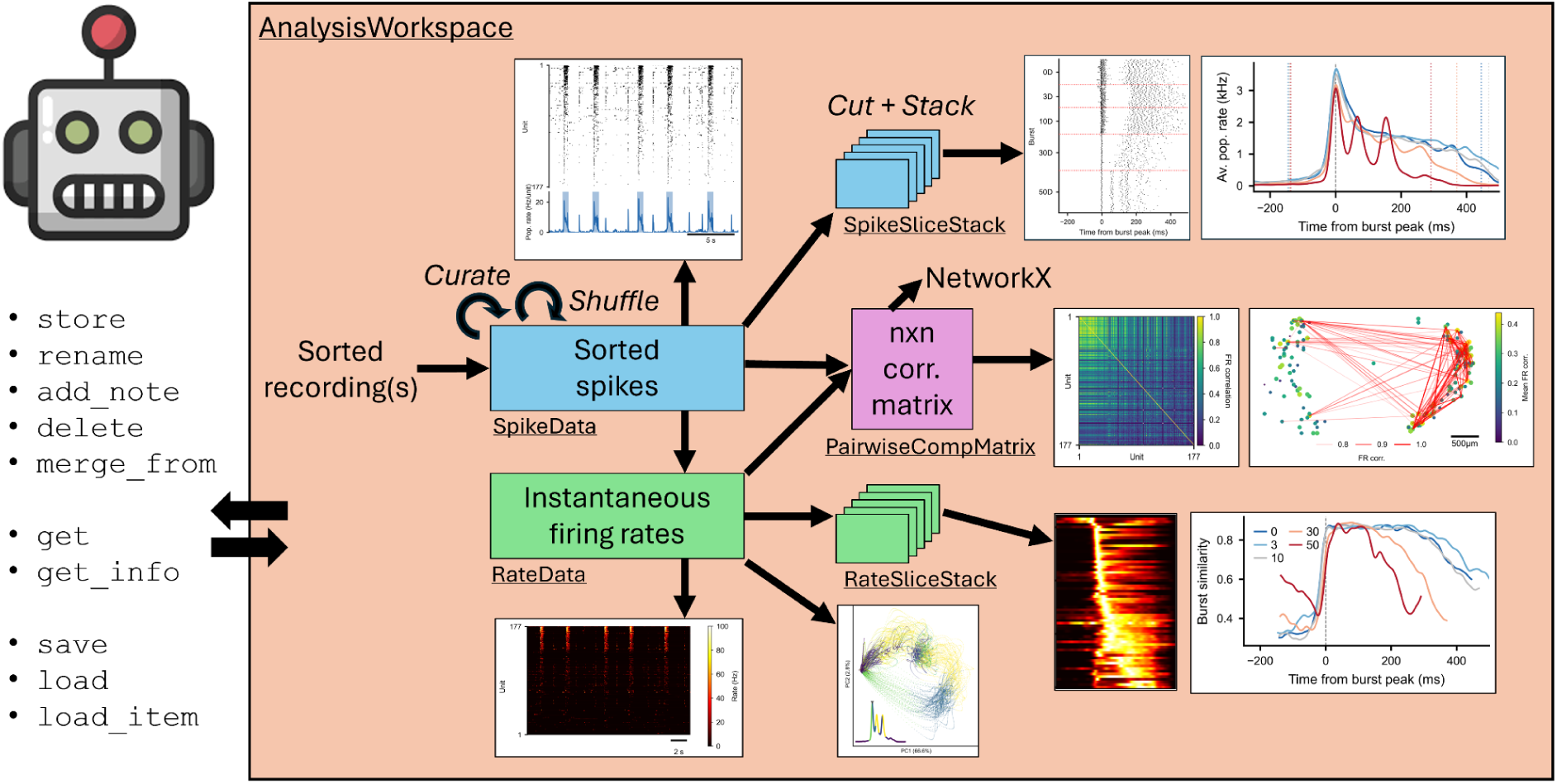
SpikeLab library architecture and data class hierarchy. The SpikeLab framework centers on a composable set of data structures for neural spike train analysis, managed through an HDF5-backed AnalysisWorkspace. Spike-sorted recordings are loaded as SpikeData objects, which can be curated (unit filtering, quality selection) and shuffled to produce surrogate datasets. From SpikeData, three derived representations are constructed: (1) RateData, a units-by-time instantaneous firing rate matrix (bottom); (2) SpikeSliceStack and RateSliceStack, collections of aligned SpikeData or RateData segments produced by event-aligned slicing, sliding-window framing, or surrogate generation (top and bottom right); and (3) PairwiseCompMatrix, an *n* x *n* comparison matrix (e.g., spike time tiling coefficients, firing rate correlations) that supports network graph construction via NetworkX (center right). Both SpikeData and RateData can generate slice stacks and pairwise comparison matrices. Each data class provides its own dedicated analysis and visualization methods (example outputs shown for each class). This modular structure forms a foundation for incorporating new analysis methods in the future. All data classes support the same temporal slicing, unit subsetting, and workspace persistence operations. The AnalysisWorkspace (left) provides caching, annotation, and retrieval of all intermediate results, enabling reproducible analysis across sessions.

SpikeData is the central representation: it stores spike trains for an arbitrary number of units alongside neuron-level metadata, and provides methods for computing firing rates, detecting population bursts, estimating inter-spike interval statistics^26^, and generating population rate traces.^27^ SpikeData objects can be curated (for example filtering units based on cluster quality metrics or by brain region where they were recorded), sliced in time (‘subtime()’), subsetted by unit (‘subset()’), and shuffled to produce surrogate datasets, all returning new SpikeData objects that are immediately compatible with every downstream method.

RateData stores instantaneous firing rate matrices (units x time) derived from SpikeData through various estimation methods (e.g., binned spike counts, Gaussian convolution, kernel-smoothed ISI.^28,29^ RateData supports the same temporal slicing and unit subsetting operations, and serves as the input for rate-based analyses such as pairwise firing rate correlations and dimensionality reduction.

SpikeSliceStack and RateSliceStack are collections of aligned SpikeData or RateData segments, respectively. They are constructed through three main routes: (1) event-aligned slicing (‘align_to_events()’), which aligns activity to external or internal events; (2) sliding-window framing (‘frames()’), which partitions a recording into overlapping temporal windows; and (3) surrogate generation (‘spike_shuffle_stack()’), which produces degree-preserving shuffled datasets for statistical testing. Because each slice in a stack is a full SpikeData or RateData object, any analytical operation available on a single recording can be applied uniformly across all slices in the stack.

PairwiseCompMatrix stores an *n* x *n* comparison matrix (e.g., spike time tiling coefficients (STTC)^30^ or firing rate correlations^29,31^ together with metadata describing the comparison method and parameters. A PairwiseCompMatrixStack collects multiple such matrices. For example, one per experimental condition or one per slice in a stack, enabling direct cross-condition comparison of pairwise structure. Both classes provide methods for visualization and matrix manipulation (e.g., thresholding), as well as NetworkX exports for graph-based analyses.^32^

All data classes can be persisted through the AnalysisWorkspace, an HDF5-backed storage layer that can hold any number of recordings alongside the results of their analyses and cross-recording comparisons, with support for annotation with metadata and retrieval across analysis sessions. The workspace ensures reproducibility: once a computation is stored, re-running a script loads the cached result rather than recomputing it.

### Agentic interfaces

SpikeLab provides two complementary interfaces for agentic access to the library: (1) a skill-based system for interactive analysis, and (2) an MCP server for programmatic access.

### Skill system

The skill system is the primary agentic interface. Users interact with SpikeLab through a single entry-point skill (/spikelab) that handles library installation and environment setup. This skill automatically routes requests to the appropriate specialized role based on intent. Four such roles are available.

The analysis-implementer skill translates natural language analysis requests into executable code using the library’s API. Its behavior is governed by a set of explicit constraints that enforce bounded autonomy:

- File scope: The skill may only create or modify files within the user’s analysis directory. It cannot modify library source code, ensuring that the underlying methods remain stable across sessions.
- Mandatory library usage: All neuroscience-specific computations (e.g. spike correlations, burst detection, population coupling, firing rate estimation) must use SpikeLab library methods. The skill is prohibited from implementing custom neuroscience logic independently, preventing *ad hoc* method invention and ensuring that analyses follow established, peer-reviewed methodologies with reproducible results.
- Correctness over efficiency: The skill must faithfully execute the user’s request without silently reducing data windows, downsampling, skipping units, or coarsening temporal resolution. If a computation is genuinely intractable, it must warn the user and propose alternatives rather than applying hidden shortcuts.
- Clarification-seeking: Scientific decisions such as how to operationalize experimental conditions, which time windows to analyze, or which method to apply remain with the researcher. When a request is ambiguous, the skill asks for clarification before proceeding.
- Workspace caching: Expensive intermediate results are cached in the AnalysisWorkspace. The skill checks for cached results before recomputing and never overwrites existing workspace keys without explicit user permission.
- Analysis logging: The skill maintains an analysis log documenting experiment context, methods, findings, and open questions, updated at the end of every session. This produces an audit trail of analytical decisions without the user needing to request it.

The educator skill provides a read-only complement. The educator explains concepts, interprets results, and answers questions about the library’s methods and the underlying neuroscience.

The educator never creates, modifies, or executes any files. If a user asks the educator to run an analysis, it redirects to the analysis-implementer. This separation ensures that explanations are never conflated with execution, and that the educator cannot inadvertently modify data or results.

The spike-sorter skill manages an optional upstream pipeline that can produce sorted recordings which in turn can be consumed by the analysis-implementer. This module is largely built on top of SpikeInterface^24^, it runs and configures spike sorting algorithms (e.g., Kilosort^25^), applies automated and interactive curation steps, and outputs curated SpikeData objects with full provenance metadata ready for analysis. Like the analysis-implementer, it operates under bounded file scope and mandatory library usage constraints (more information in Supplementary Information: Spike sorting integration). Alternatively, SpikeLab also provides expansive tooling for importing spike sorted data directly.

The developer skill closes the loop between analysis and library development. When an analysis script contains novel computations that are general enough to be reusable, the developer skill integrates them into the library. The developer audits the script against existing methods, replaces reimplementations with library calls, adapts novel computations to library conventions, writes tests covering main usage and edge cases, and exposes new methods through the MCP server where appropriate. This enables a cycle where analyses conducted through the implementer can feed back into the library itself, expanding its capabilities over time.

For computationally intensive tasks, the implementer and spike-sorter skills can submit jobs to remote Kubernetes clusters via an optional batch-jobs module (see Supplementary Materials: Batch-jobs).

Together, the four skills define a workflow covering the full path from raw recordings to interpreted results and back to library development. The spike-sorter produces curated data, the implementer translates natural language requests into validated analysis code, the educator provides interpretation and context, and the developer integrates proven analyses back into the library, all without the researcher needing to interact with the code directly.

### MCP server

As an alternative to the skill-based workflow, SpikeLab exposes its full analytical capabilities through a Model Context Protocol (MCP) server.^36^ The MCP server wraps the library’s data classes and methods as individually callable tools, providing stateless programmatic access suitable for integration into automated pipelines or external agent frameworks. Where the skill system operates within a conversational context with workspace persistence and analysis logging, the MCP server offers a lightweight, tool-by-tool interface that can be composed freely by any MCP-compatible client.

The following sections evaluate this architecture empirically, first through a controlled benchmark against unassisted frontier models and then through its application to diverse recording modalities and a multi-condition pharmacological study.

### Agentic tools improve analysis quality and reproducibility

To test whether domain-specific agentic tools improve the quality of LLM-driven neuroscience analysis, we designed a benchmark comparing three conditions on a standardized four-task analysis pipeline: (1) Claude Opus 4.6 without library access (“Opus” or O), (2) Claude Sonnet 4.6 without library access (“Sonnet” or S), and (3) Claude Sonnet 4.6 with SpikeLab’s analysis-implementer skill (“SpikeLab” or SL). At the time of benchmarking (March 24th-26th 2026), Opus 4.6 and Sonnet 4.6 were Anthropic’s most and second most powerful models, respectively. Opus was run at high reasoning effort while both Sonnet and SpikeLab conditions used medium effort. Each condition was run three times on the same prompts, allowing us to assess both correctness and reproducibility. Opus and Sonnet started with a bare conda environment (Python 3.10) and were responsible for installing any packages they needed, including standard scientific libraries such as numpy, scipy, and matplotlib, but had no access to domain-specific tools. The SpikeLab condition started with the same bare environment plus a pre-installed SpikeLab package with all its dependencies. All conditions received identical task prompts (**Fig. 2A**), with the sole exception that the SpikeLab condition was prefixed with the skill invocation command on Task 1.

**Figure 2.**
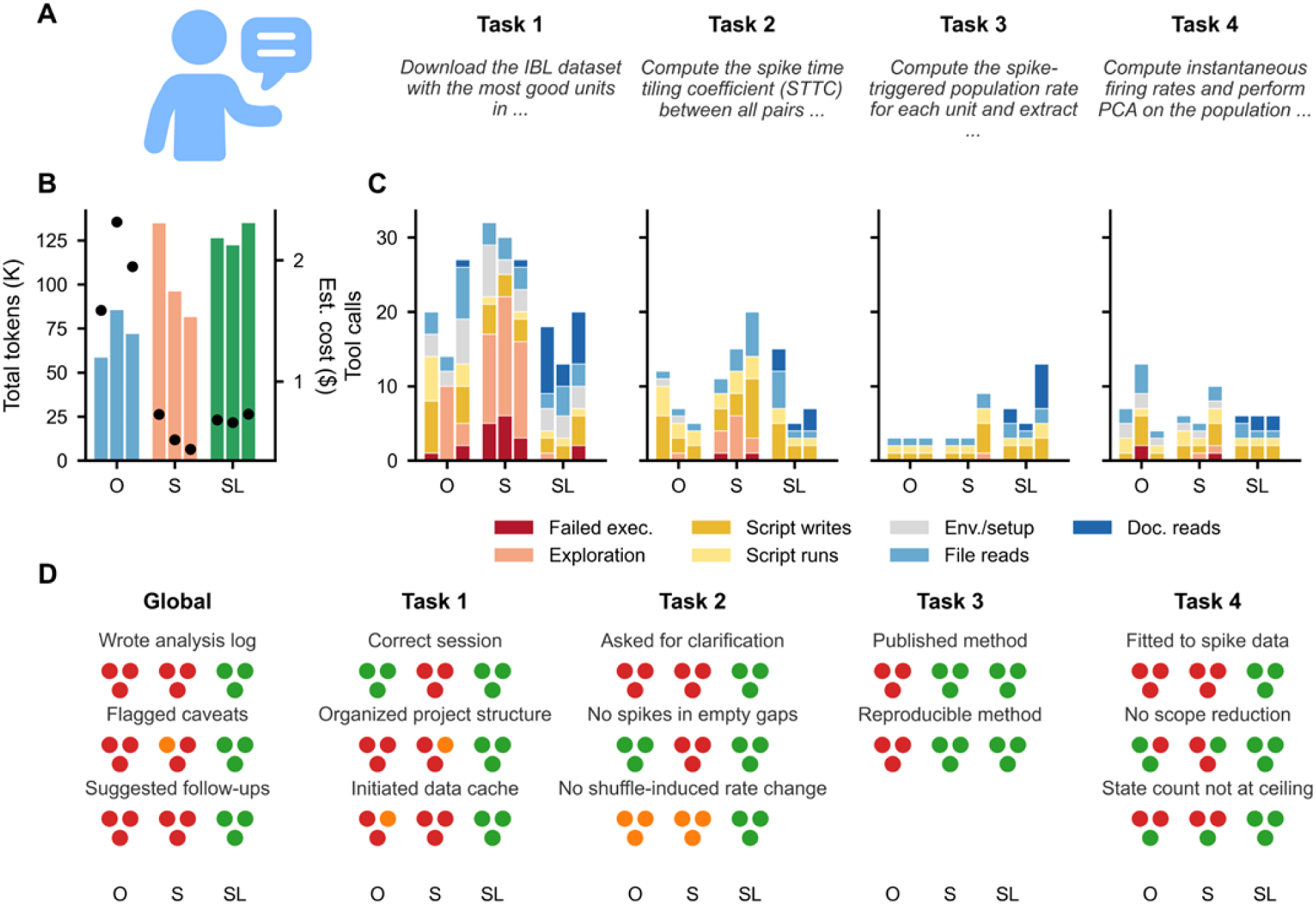
**Benchmarking SpikeLab analysis tools against frontier models.** (**A**) Prompts provided to each model condition. All conditions received identical prompts except that the SpikeLab condition was prefixed with “with /spikelab,” on Task 1. Each task builds on the same IBL auditory cortex dataset loaded in Task 1. The Task 2 prompt was deliberately underspecified (e.g., “cue and no-cue parts”) to test whether models would ask for clarification or make assumptions. See *Methods: Benchmark protocols* for the verbatim prompts per task. (**B**) Total tokens consumed across all four tasks per run (3 runs per condition). Bars show token count (left y-axis, in thousands). Circles indicate estimated API cost (right y-axis), calculated using a blended rate assuming 80% input / 20% output token split (Opus: $27/M tokens; Sonnet: $5.4/M tokens). Despite using fewer tokens per session, Opus’s higher per-token cost results in a greater total cost. O = Opus, S = Sonnet, SL = Sonnet + SpikeLab. (**C**) Stacked bar charts showing the composition of tool calls per run per task, decomposed into seven categories. Failed executions (dark red) and inline exploration (light orange) represent trial-and-error behavior: probing APIs, debugging errors, and running one-off commands. Script writes (amber) and successful script runs (yellow) represent productive code development. Environment/setup (light grey) covers package installation, directory creation, and skill loading. File reads (light blue) and documentation reads (dark blue) represent information gathering, with documentation reads occurring almost exclusively in the SpikeLab condition. Each group of bars within a task subplot represents one condition (O, S, SL), with individual bars for each of the 3 runs. (**D**) Per-run assessment of key methodological and workflow issues across tasks. Each triangle of three circles represents one condition (O = Opus, S = Sonnet, SL = Sonnet + SpikeLab), with individual circles for each of the 3 runs. Green = correct/present, orange = partially correct, red = incorrect/absent.

The four tasks represent a progressive neuroscience analysis workflow on publicly available data from the International Brain Laboratory (IBL): (1) identify and load the IBL probe with the most good-quality units in auditory cortex, (2) compute spike time tiling coefficients with shuffle-based z-scores, separately for stimulus-driven and spontaneous periods, (3) quantify population coupling for each neuron, and (4) fit a latent state-space model to the population activity (see *Methods: Benchmark protocol* for verbatim prompts for each task). Task 2 was deliberately underspecified. The prompt “run this analysis separately on the cue and no-cue parts of the recording” admits multiple valid interpretations to test whether models would seek clarification or proceed with an unexamined assumption.

### Resource usage: tokens, cost, and tool call composition

Opus consumed the fewest tokens across all three runs (58,780–85,720 per session, mean 72,202) compared to Sonnet (81,622–135,010, mean 104,288) and SpikeLab (122,370–135,131, mean 127,999) (Fig. 2B). However, Opus’s 5x higher per-token cost makes it the most expensive condition despite its lower token usage, with a blended rate of $27/M tokens (80% input and 20% output). Opus sessions averaged $1.95 each, compared to $0.56 for Sonnet and $0.69 for SpikeLab at $5.4/M tokens.

Tool call counts provide a complementary view of agentic effort (**Fig. 2C**; classification rules and verification details in Methods; raw session transcripts and line-by-line evidence in Supplementary Data). Total tool calls per session averaged 39.3 for Opus (range 37–42), 57.0 for Sonnet (52–66), and 40.3 for SpikeLab (29–46). SpikeLab’s comparable total to Opus reflects a fundamentally different composition, where Opus and Sonnet spent calls on trial-and-error API exploration and failed executions. SpikeLab invested in documentation reads (1–9 per task) that informed correct implementations on the first attempt. The largest divergence between Opus and Sonnet occurred in Task 1: Opus averaged 20.3 calls, Sonnet 29.7, and SpikeLab 17.0. Tasks 2–4 showed smaller differences: Task 2 averaged 8.0 (Opus), 15.3 (Sonnet), 9.0 (SpikeLab); Task 3 averaged 3.0 (Opus), 5.0 (Sonnet), 8.3 (SpikeLab); Task 4 averaged 8.0 (Opus), 7.0 (Sonnet), 6.0 (SpikeLab). SpikeLab’s higher Task 3 count (8.3 vs 3.0–5.0) reflects one run where the agent discovered a bug mid-task and performed additional documentation reads to diagnose and fix it. We report tool call counts rather than wall-clock time because runtime is confounded by the models’ computational choices. For example, fitting an HMM on the full recording versus a downsampled subset, or using a Numba-accelerated STTC implementation versus pure Python.

The composition of tool calls reveals qualitatively different working strategies (**Fig. 2C**). The largest divergence occurred in Task 1 (data discovery and loading), where both Opus and Sonnet engaged in extensive trial-and-error exploration of the IBL API. Sonnet made 12–16 inline‘python-c’ calls per Task 1 run, one-liner probes testing API endpoints, checking data shapes, and inspecting metadata. Opus used fewer inline probes (0–10) but wrote more throwaway scripts (up to 7 per run). Sonnet produced substantial failed executions: 3–6 per run in Task 1 alone; Opus had 0–2. SpikeLab replaced this exploration with documentation reading, performing 3–9 reads of the library’s repo maps and API reference during Task 1, and completed the task with only 2 failed executions across all three runs combined (both minor attribute errors in a single run, not methodological bugs). Once the documentation was read in Task 1, it remained in the conversation context, reducing the need for additional reads in subsequent tasks (1–6 reads per task for Tasks 2–4, compared to 3–9 for Task 1). Beyond Task 1, the main distinction was consistency. SpikeLab’s per-task tool counts varied narrowly across runs (e.g., Task 4: 6, 6, 6), while Opus and Sonnet showed wider run-to-run variation reflecting different debugging paths and parameter choices (e.g., Opus Task 4: 7, 13, 4; Sonnet Task 2: 11, 15, 20).

Next, we evaluated each run against a 14-issue scorecard (**Fig. 2D**) spanning data discovery, methodological correctness, reproducibility, and scientific documentation practices. Each issue was scored as correct (green), partially correct (orange), or incorrect/absent (red) for each of the nine runs. Detailed code examples for each issue are provided in Supplementary Materials; raw session transcripts and all generated analysis scripts and figures are available in Supplementary Data. The scorecard reveals systematic differences between conditions that are not attributable to stochastic variation. For example, the same model hits the same failure modes in every run, while SpikeLab-assisted runs avoid them entirely.

### Data discovery: API strategy determines downstream power

The first task, finding the best IBL probe, produced a striking divergence. All three Opus runs correctly identified the probe insertion with the most good-quality auditory cortex units: 208 units from session‘7f5df7eb’, probe‘0c15a331’. All three Sonnet runs found a different, inferior session with only 69 auditory cortex units (‘ff48aa1d’). All three SpikeLab runs also found the correct 208-unit session. Each model was deterministic and locked into a single API strategy on its first attempt and never reconsidered.

The root cause was an undocumented gap between two IBL API endpoints:‘one.search()’ queries session-level metadata tags (which were incomplete for this session), while‘one.search_insertions()’ queries probe trajectory data. Sonnet used the former, Opus happened to discover the latter. Neither model flagged the endpoint selection as a methodological choice that could affect results. SpikeLab’s‘query_ibl_probes()’ sidesteps this class of problem entirely by downloading the complete Brain-Wide Map unit table containing every unit from every probe across the entire IBL database, making it impossible to miss a session regardless of how it is indexed.

### Project organization and data caching

Upon accessing the IBL dataset, the three conditions diverged in how they organized their work. Opus and Sonnet produced flat directories with 4–11 files including throwaway scripts (e.g.,‘find_session.py’ through‘find_session5.py’); SpikeLab created structured directories with separate subdirectories for figures, results, and cache from the start, with no throwaway files.

Opus and Sonnet re-downloaded data from the IBL database for nearly every task script, 21 of 24 final scripts across all runs called the ONE API directly. SpikeLab cached downloaded data to serialize using the Python pickle module on the first task and stored intermediate results in per-task HDF5 workspace files, so re-running any script skips computation entirely.

### Ambiguity handling: clarification-seeking as a quality signal

In the Task 2 prompt, “run this analysis separately on the cue and no-cue parts of the recording” was deliberately ambiguous. “No-cue” could refer to inter-trial intervals, catch trials, or pre/post-task periods, each corresponding to a different experimental comparison and a different scientific question. Neither Opus nor Sonnet asked for clarification in any of the six runs. Opus consistently interpreted “no-cue” as inter-trial intervals; Sonnet switched between trial-type contrasts and an ITI-based approach across runs, producing two fundamentally different experimental designs without the prompt changing.

SpikeLab asked for clarification in all three runs, presenting explicit interpretations before proceeding. This behavior is a direct consequence of the analysis-implementer skill’s “never assume, ask if unsure” rule. The skill converts a general LLM tendency to proceed with a default

assumption into a concrete clarification step, ensuring that the experimental design reflects the researcher’s intent rather than the model’s arbitrary interpretation.

### Shuffle procedures: deterministic bugs in Opus and Sonnet

Task 2 required computing STTC with shuffle-based z-scores to identify significantly correlated neuron pairs. Both Opus and Sonnet encountered bugs in their shuffle implementations and each model hit the same bug class in every run, revealing systematic failure modes rather than stochastic errors.

Opus concatenated cue intervals into a single continuous stream and applied circular shifts within it. This destroys trial-to-trial firing rate variability (e.g., differences between correct and incorrect trials), flattening the structure that the shuffle null should preserve. Sonnet concatenated cue intervals with 40 ms gaps between them and applied circular shifts. The shift can cause spikes to land in these empty gap zones. Moreover, trial-to-trial firing rate variability is destroyed as with Opus. Both models’ circular shift approaches preserved the overall average firing rate per unit and so neither fully invalidated the null distribution. Still the finer temporal structure that a well-designed shuffle should maintain was lost.

Sonnet also encountered a numerical edge case scenario for low-firing-rate neuron pairs. The 10 shuffled STTC values were nearly identical (sigma approaching 0), producing extreme z-scores (z > 500). The fix applied arbitrary clamping rather than flagging unreliable pairs, silently distorting the z-score distribution. The clamping threshold itself varied across runs, adding another source of cross-run inconsistency.

SpikeLab avoided these issues entirely by using‘spike_shuffle_stack()’, a degree-preserving shuffle method that operates on the binary raster matrix^27^, exactly preserving both per-unit spike counts and per-time bin population rates. Where shuffle variance was near zero, the z-score was set to NaN rather than clamped, an explicit signal of unreliable estimation rather than a silent distortion.

It is important to note that both Opus’s and Sonnet’s bugs were deterministic here. The same model produced the same bug class in every run. These are not random errors that might self-correct with repeated attempts. Instead, they are systematic failure modes arising from the model’s default approach to a domain-specific problem. A library method that encodes the correct procedure eliminates this issue.

### Population coupling: method invention versus method application

Task 3 asked each condition to compute population coupling, the degree to which individual neurons’ firing is linked to overall network activity. SpikeLab’s library implements the spike-triggered population rate (stPR) method^38^, building on previous implementations^39^ with added baseline subtraction and firing-rate normalization.

Sonnet implemented a method equivalent to these previous implementations^39^, consistent across all three runs but lacking baseline subtraction and normalization. Opus invented an ad-hoc approach from first principles each time, producing three fundamentally different metrics in different units across three runs: a dimensionless z-score, a unitless ratio, and a raw firing rate in spikes per second. These quantities cannot be compared across runs. The model reinvented a different metric each time because there was no stable reference implementation to anchor it.

SpikeLab called a single library function (‘compute_spike_trig_pop_rate()’) with default parameters, encoding the full Bimbard method: leave-one-out population rate, baseline subtraction, normalization by the sum of mean firing rates, Butterworth low-pass filtering at 20 Hz, and a +-80 ms lag curve. Across all three runs, the coupling values were identical: range-0.14 to +0.43, median 0.066, with the same unit (index 35, AUDd, 1.85 Hz) selected as closest to the median every time.

Task 3 was the “easiest” in terms of execution. All conditions produced working code on the first attempt with no runtime errors. But the absence of implementation bugs does not imply methodological correctness. Without a reference implementation encoding the published method, even a capable model defaults to *ad hoc* approaches that may be reasonable in isolation but are not stable, comparable, or grounded in the literature.

### Latent state models: scope reduction and model selection

Task 4 asked each condition to fit a latent state-space model. Both Opus and Sonnet independently converged on the same pipeline: bin, smooth, z-score, PCA, fit GaussianHMM, select *K* by BIC. This suggests that this pipeline is a well-represented recipe in training data. SpikeLab used a recent Gaussian Process Latent Variable Model (GPLVM) as introduced by Zheng et al., which determines effective dimensionality from the data via the GP prior rather than requiring discrete state count selection.^40^ Either approach is a valid interpretation of the prompt. However, the convergence on PCA+HMM illustrates that models default to well-established methods represented in their training data, while more recent approaches like GPLVM are unlikely to surface without domain-specific tooling that makes them accessible.

The prompt asked to “fit a latent state space model to the spike data,” but both Opus and Sonnet fitted the HMM to a PCA-projected representation rather than to the spike data directly. In addition, both silently coarsened the temporal resolution in a majority of runs to reduce computation time: Opus rebinned from 50 ms to 100 ms and subsampled every 3rd bin (300 ms effective resolution) in one run; Similarly, Sonnet kept only every 2nd or every 4th time bin before fitting the HMM in two of three runs. The reduced temporal resolution limits the model’s ability to resolve fast state transitions. SpikeLab used the full 77.6-minute recording (93,085 bins at 50 ms) in all three runs, consistent with the analysis-implementer skill’s “correctness over efficiency” mandate.

Additionally, both models hit the upper bound of their HMM state search range in 4 of 6 runs (Opus: *K*=8 in runs 1–2; Sonnet: *K*=6–7 in runs 1–2) without applying the standard sanity check to see whether BIC was still decreasing at the boundary. Neither extended the search range.

GPLVM sidesteps this entirely: it used 100 latent bins, with the posterior showing near-uniform occupancy across visited states.

Crucially, none of these decisions: the PCA projection, the temporal downsampling, or the unchecked BIC boundary, were reported to the user. The models cut corners to make the computation tractable and made methodological choices without flagging them, illustrating the risk of leaving too much autonomy to models operating without domain-specific guardrails.

### Scientific practices: documentation, caveats, and follow-ups

Beyond task-specific findings, the benchmark revealed systematic differences in scientific workflow practices (**Fig. 2D**, Global category). Only SpikeLab created analysis documentation. In all three runs, an‘ANALYSIS_LOG.md’ file was created and updated after each task, unprompted, recording experimental context, methods, findings, and open questions. Neither Opus nor Sonnet wrote any documentation in any of their six runs. SpikeLab also proactively flagged caveats in all three runs (e.g., ITI duration imbalance affecting z-score reliability) and suggested follow-up analyses (e.g., separate AUDd vs. AUDv comparison). Opus never flagged caveats; Sonnet did so inconsistently, noting z-score clamping in one run but not the other two.

### Summary

The benchmark reveals a clear hierarchy. SpikeLab-assisted analysis was correct and reproducible across all three runs for all four tasks. Sonnet used consistent implementations for most tasks but switched between fundamentally different experimental designs for the cue/no-cue comparison across runs and applied silent workarounds for numerical edge cases. Opus demonstrated strong initial problem-solving (finding the correct session, catching bugs mid analysis) but suffered from method drift through inventing a different coupling formula or a different plotting approach in each run. A recurring pattern across both Opus and Sonnet is silent decision-making. Neither model clearly reported to the user that methodological choices were being made: the API endpoint selection (Task 1), the cue/no-cue operationalization (Task 2), the coupling formula (Task 3), and the temporal downsampling and PCA projection (Task 4) were all implemented without flagging them as decisions that could affect results. Without a library encoding domain-specific methods and a skill system that mandates transparency, even frontier models default to *ad hoc* solutions that vary across invocations and go unreported.

These findings demonstrate that overly autonomous model-driven analysis can introduce silent methodological errors, unreported decisions, and cross-run inconsistencies that, if left unchecked, risk undermining the reliability of scientific results. Yet the models themselves demonstrated strong capabilities, which include exploring unfamiliar APIs, writing working analysis pipelines from scratch, and catching their own errors. The limitation is not reasoning ability but the absence of stable, validated building blocks. SpikeLab provides these building blocks in a way that is rigid where correctness matters (shuffle methods, published coupling formulas, full-resolution analysis) and flexible where the models excel (composing analyses, interpreting results, adapting to new datasets). The combination leverages the strengths of both, pairing the speed and versatility of frontier models with the methodological reliability of domain expertise embedded into reusable library methods and agentic skill instructions. Having

established that agentic tooling improves analysis quality on a single dataset, we next showcase how the same pipeline generalizes across recording modalities and species.

### A unified pipeline across recording modalities

All analyses presented in the following sections were conducted fully through SpikeLab’s text-to-analysis workflow, meaning that not a single line of code was written manually. These examples demonstrate that the agentic pipeline generalizes across recording modalities and experimental paradigms without requiring the user to write or modify analysis code directly. We applied the library to three electrophysiology preparations that differ in species, recording technology, and experimental context: (1) mouse auditory cortex recorded *in vivo* with Neuropixels probes^33^, (2) human anterior temporal lobe recorded *in vivo* with dual 64-channel Utah arrays^34^, and (3) a human forebrain organoid recorded *in vitro* on a Maxwell Biosystems MaxOne high-density microelectrode array.^35^ All three spike-sorted datasets were loaded as SpikeData objects and analyzed through the same pipeline of library methods without modification to the analysis code.

Despite qualitatively different activity patterns, including dense spontaneous firing in mouse cortex (**Fig. 3A**), sparser irregular activity in human cortex (**Fig. 3B**), and synchronized population bursts in the organoid (**Fig. 3C**), all three recordings are represented as the same SpikeData object and can be processed through the same analysis and visualization methods.

**Figure 3.**
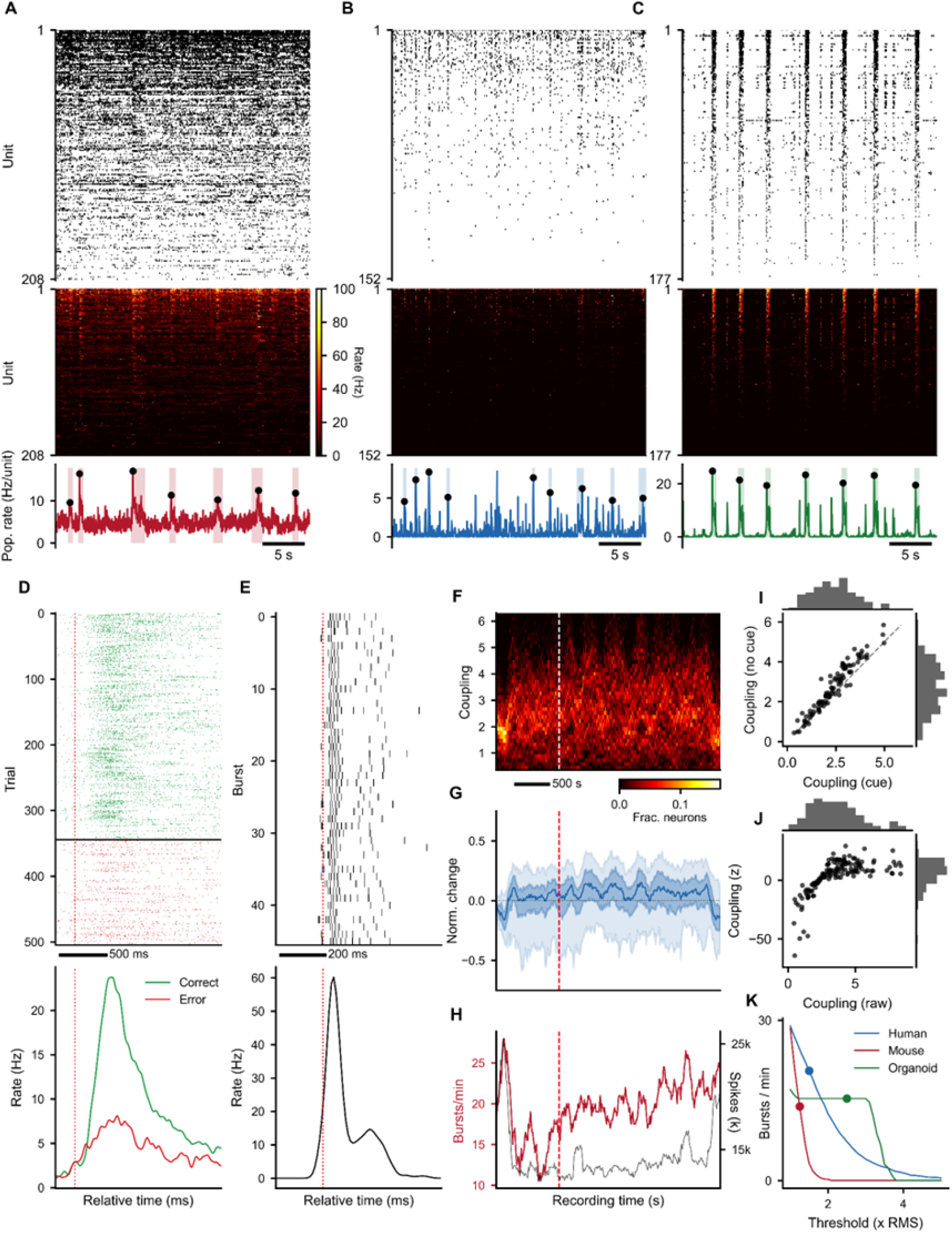
SpikeLab applied to diverse electrophysiology preparations, mouse auditory cortex, human anterior temporal lobe, and human forebrain organoid. (**A**) Representative 30 s recording from mouse auditory cortex (IBL dataset, eid: 7f5df7eb, pid: 0c15a331, 208 AUDv/AUDd units). Top: spike raster sorted by firing rate. Middle: instantaneous firing rate heatmap. Bottom: population rate with detected burst windows (shaded). (**B**) Same as (**A**) for human anterior temporal lobe. (**C**) Same as (**A**) for a human forebrain organoid. (**D**) Single-unit stimulus-aligned raster (top) and Gaussian-smoothed peri-stimulus time histogram (bottom) for an example mouse auditory cortex unit (unit 36). Trials split by correct (green) and error (red) outcomes. Dashed red line: stimulus onset. (**E**) Single-unit burst-aligned raster (top) and Gaussian-smoothed peri-event time histogram (bottom) for an example organoid unit (unit 85). Dashed red line: burst peak. (**F**) Population coupling distribution over the full human recording (cue + no-cue periods). Heatmap shows the fraction of neurons at each coupling value across sliding 2-minute windows (10 s step). Dashed white line: end of cue period. (**G**) Normalized change in population coupling relative to the first window, shown as percentile bands (5th-95th and 25th-75th percentiles). Normalization: *N* = (x - x_0_) / (x + x_0_), where x_0_ is the first-window value; negative values indicate decreased coupling, positive values indicate increased coupling. Dashed red line: end of cue period. (**H**) Burst rate (red, left axis) and total spike count (black, right axis) per 2-minute sliding window across the full recording. Dashed red line: end of cue period. (**I**) Mean population coupling during the cue period vs. the no-cue period per unit. Dashed line: identity. (**J**) Raw population coupling vs. shuffle-normalized (z-scored) population coupling per unit (*n* = 151 units). Z-scores computed against 100 degree-preserving spike shuffles. (**K**) Burst detection sensitivity: burst rate as a function of the population rate threshold (RMS multiplier) for each preparation. Dots indicate the threshold used for each recording.

### Diverse SliceStack applications: event alignment, framing, and shuffling

A central design element of SpikeLab is the SliceStack, a collection of aligned SpikeData or RateData segments that enables event-locked, windowed, and surrogate-based analyses through a single interface. Here we illustrate four distinct uses of this abstraction.

- Stimulus-aligned slicing: In the mouse recording, trials were aligned to stimulus onset and split by behavioral outcome (**Fig. 3D**). Example unit 36 shows a transient increase in firing rate following stimulus presentation, with a clear separation between correct and error trials. This analysis used‘align_to_events()’ with externally defined stimulus times to construct a SpikeSliceStack. The same approach generalizes to any externally timed event such as electrical or optical stimulation, pharmacological application onset or behavioral markers.
- Activity-aligned slicing: In the organoid, where no external stimulus is present, we aligned activity to internally generated population bursts detected from the population rate^29^ (**Fig. 3E**). Example unit 85 shows a sharp increase in firing rate locked to burst onset. The same‘align_to_events()’ method was used, but with burst peak times derived from the recording itself rather than from an external event log, further demonstrating that the SliceStack interface is agnostic to the source of alignment events.
- Sliding-window framing: To track how population coupling evolves over the course of the human recording, we used ‘frames()’ to partition the full recording into a stack of overlapping 2-minute SpikeData objects (10-second step) bundled into a SpikeSliceStack. Because each slice in the stack is itself a full SpikeData object, the same analytical operations can be applied uniformly across all frames. Here, we computed zero-lag spike-triggered population rate coupling per unit within each frame. The resulting time-by-unit coupling matrix shows a subtle shift toward higher coupling in the no-cue period (**Fig. 3F-G**), alongside non-monotonic changes in burst rate and spike count across the recording (**Fig. 3H**). Per-unit mean coupling confirms this shift, with especially the already strongly coupled units showing slightly further increased coupling during the no-cue period compared to the cue period (**Fig. 3I**).
- Shuffle-based normalization: To distinguish genuine coupling from coupling expected by chance given each unit’s firing rate, we applied‘spike_shuffle_stack()’ to generate 100 degree-preserving surrogate datasets and computed coupling z-scores against the shuffle distribution (**Fig. 3J**). Raw and z-scored coupling values are broadly correlated, confirming that the coupling estimates are not artifacts of firing rate alone, while the z-score normalization reveals which units are coupled beyond chance expectation.

### Cross-recording comparison

Because all three preparations share the same SpikeData representation, cross-recording analyses require no format conversion or adapter code. We illustrate this with a burst detection sensitivity analysis (**Fig. 3K**), where burst rate is plotted as a function of the population rate threshold (RMS multiplier) for all three recordings in a single panel. The resulting curves characterize the burstiness of each preparation. Here the organoid shows an initial plateau followed by a steep drop-off in burst rate with increasing threshold, reflecting frequent high-amplitude synchronized events. Meanwhile, the mouse and human recordings drop off more smoothly, indicating lower-amplitude population activity. This analysis loaded three SpikeData objects from the same workspace, iterated over a shared threshold range using the same ‘get_bursts()’ call, and combined the results, demonstrating that the unified data representation enables direct cross-recording comparison within a single analysis session. We next demonstrate how these building blocks compose into a multi-level analysis pipeline through a pharmacological dose-response study on the organoid recording.

### Case study: diazepam dose-response in a human brain organoid

To illustrate how SpikeLab’s data structures compose into a multi-level analysis pipeline across multiple recordings, we applied the library to a pharmacological dose-response experiment.^34^ The data consists of a human forebrain organoid recorded on a MaxOne high-density microelectrode array across five diazepam concentrations (0, 3, 10, 30, and 50 μM), with 177 units matched across all conditions. Each of the four figures in this section maps primarily to one of the core data classes, demonstrating how the same recording can be progressively analyzed from single-unit properties through pairwise network structure to population-level dynamics.

### Single-unit and burst-level properties

Starting from the SpikeData representation, we extracted per-unit firing statistics across all five conditions. Representative recording windows show a visible shift in activity patterns with increasing diazepam concentration, with higher diazepam concentrations producing more frequent but shorter bursts and reduced inter-burst firing (**Fig. 4A**). Per-unit distributions of firing rate, ISI coefficient of variation, population coupling, and burst participation metrics capture the heterogeneity of single-unit responses across the population (**Fig. 4B-F**). Normalized change relative to baseline summarizes the dose-dependent trajectory for each metric across all units (**Fig. 4G-K**). Burst detection sensitivity curves (**Fig. 4L**) and burst width distributions (**Fig. 4M**) characterize how population-level burst structure changes with dose. Scatter plots comparing baseline and high-dose conditions per unit reveal which units shift their coupling and burst participation most strongly (**Fig. 4N-P**). All metrics in this figure are computed from SpikeData methods: ‘rates()’, ‘interspike_intervals()’,‘compute_spike_trig_pop_rate()’,‘get_pop_rate()’,‘get_bursts()’, ‘get_frac_active()’, and‘get_frac_spikes_in_burst()’, applied identically to each condition.

**Figure 4:**
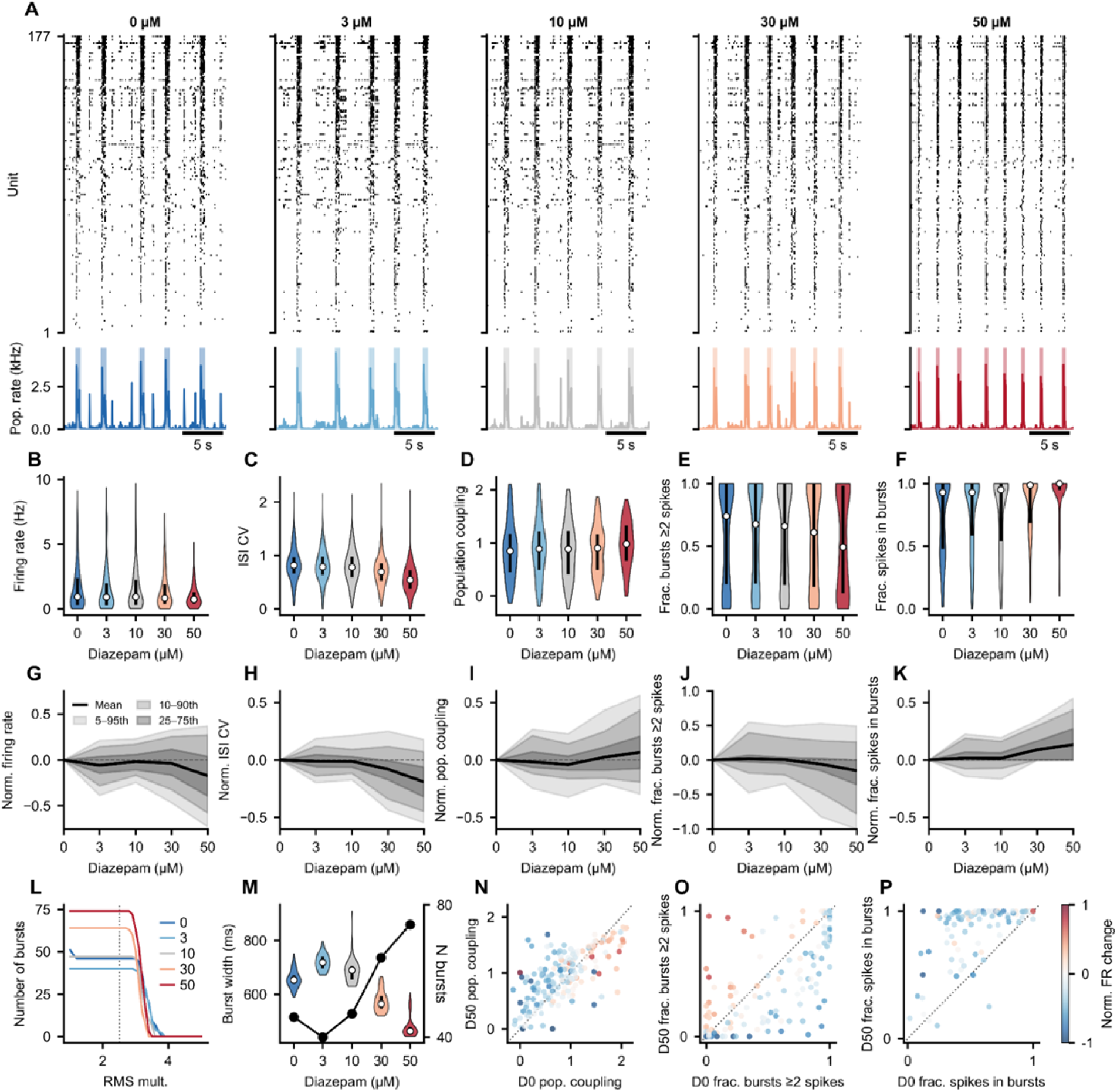
Diazepam dose-dependently alters single-unit activity and network bursting in human forebrain organoid. (**A**) Representative spike rasters (top) and population firing rates (bottom) from a 20 s window at each diazepam concentration (0, 3, 10, 30, 50 µM). Units (n = 177) are sorted by firing rate at baseline (0 µM). Shaded regions indicate detected network bursts. (**B–F**) Per-unit distributions across conditions: firing rate (**B**), ISI coefficient of variation (**C**), population coupling (**D**), fraction of bursts with ≥2 spikes (**E**), and fraction of spikes occurring within bursts (**F**). White dots indicate medians; thick black lines span the interquartile range. (**G–K**) Symmetric normalized change relative to baseline for the same five metrics, shown as percentile bands. Shaded regions indicate the 5th–95th, 10th–90th, and 25th–75th percentile ranges (light to dark); black line shows the mean. Normalization: *N* = (x’ − x₀’) / (x’ + x₀’), where x’ = max(x, 0). (**L**) Burst detection sensitivity: number of detected bursts as a function of the population rate threshold (RMS multiplier). Dashed line indicates the threshold used (2.5×). (**M**) Burst width distributions with burst counts (black line, right axis). (**N–P**) Baseline (0 µM) vs. high-dose (50 µM) scatter plots for population coupling (**N**), fraction of bursts with ≥2 spikes (O), and fraction of spikes in bursts (**P**). Points colored by normalized firing rate change. Dotted line: identity.

### Pairwise correlations and network structure

The PairwiseCompMatrix and PairwiseCompMatrixStack classes enable systematic comparison of functional connectivity across conditions. Utilizing neural spike times (**Fig. 5A**, top) and instantaneous firing rates (**Fig. 5A**, bottom), STTC and firing rate correlation matrices were computed for all 15576 unique unit pairs per condition (**Fig. 5B-C**), with their joint distribution revealing the relationship between rate-based and spike-timing-based measures of pairwise similarity (**Fig. 5D**). Per-pair correlation distributions and normalized change bands track the dose-dependent shift in connectivity strength (**Fig. 5E-F**) marking increased coherence after diazepam treatment. Network visualizations provide spatial network maps (**Fig. 5G**) and graph exports via NetworkX enable computation of graph-theoretic metrics across conditions, including mean weighted degree, clustering coefficient, path length, betweenness centrality, modularity, and rich-club coefficient (**Fig. 5H-M**). The key library methods underlying this figure are‘spike_time_tilings()’,‘get_pairwise_fr_corr()’,‘to_networkx()’, and‘plot_spatial_network()’, each applied identically across conditions. This demonstrates how a single PairwiseCompMatrixStack, holding one matrix per condition, feeds directly into both statistical summaries and graph-based network analyses.

**Figure 5:**
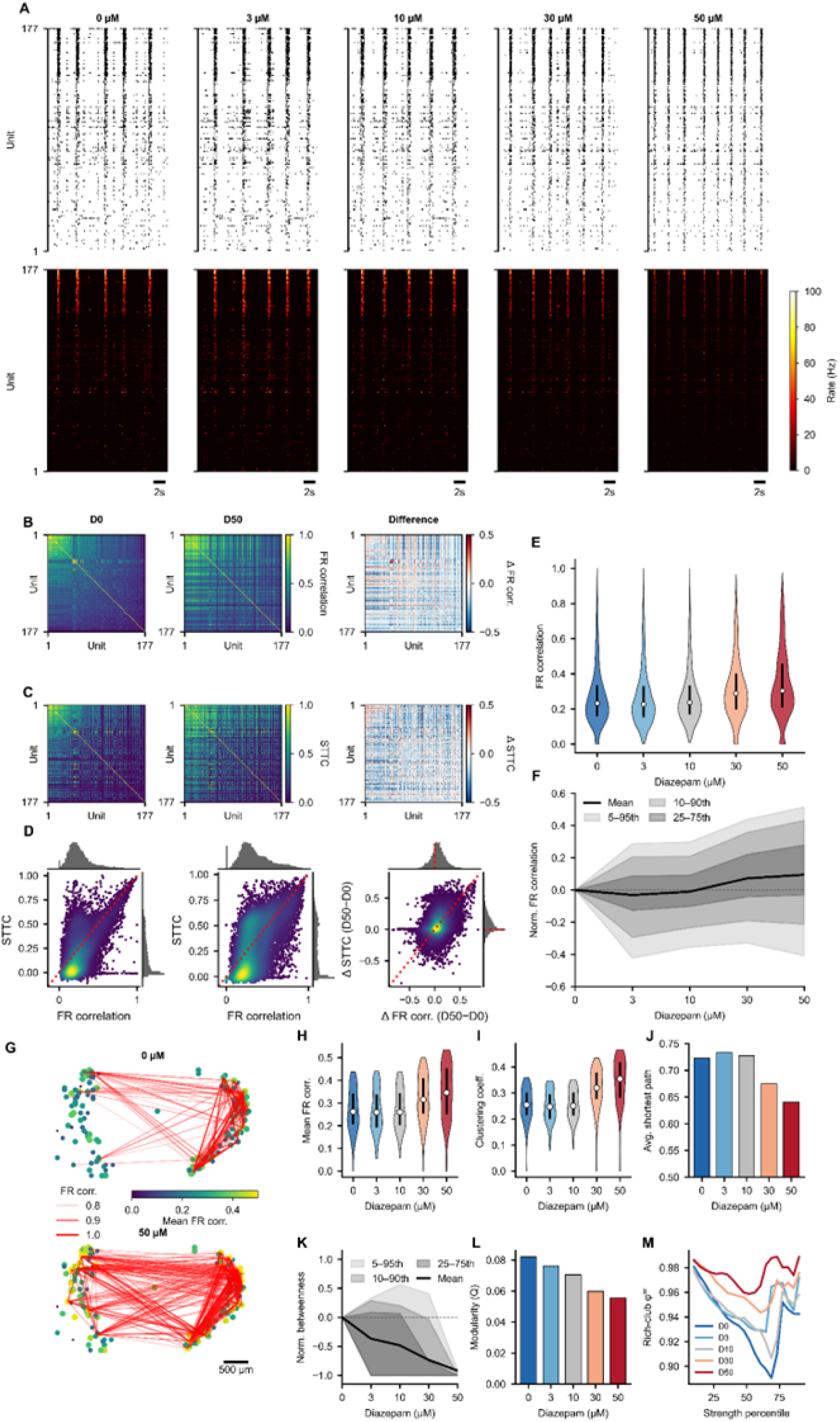
Diazepam strengthens pairwise functional connectivity. (**A**) Spike rasters (top) and instantaneous firing rate heatmaps (bottom) for each diazepam concentration (0–50 µM). Units (n = 177) sorted by mean pairwise firing rate (FR) correlation at baseline. (**B**) Pairwise FR correlation matrices at 0 µM (left), 50 µM (middle), and their difference (right). Units ordered by mean FR correlation at baseline (high to low). (**C**) Same layout as (**B**) for spike time tiling coefficient (STTC, Δ*t* = 20 ms). (**D**) FR correlation vs. STTC for all unit pairs at 0 µM (left), 50 µM (middle), and their change (right). Points colored by density; diagonal line indicates identity. Marginal histograms shown on each axis. (**E**) Per-pair FR correlation distributions across conditions. White dots: medians; thick lines: interquartile range. (**F**) Normalized FR correlation change relative to baseline, shown as percentile bands (5th–95th, 10th–90th, 25th–75th; light to dark). Black line: mean. (**G**) Spatial network maps at 0 µM (top) and 50 µM (bottom). Units plotted at MEA electrode positions, sized and colored by mean FR correlation. Red edges connect pairs with FR correlation > 0.8; edge opacity scales with correlation strength. Scale bar: 500 µm. (**H**) Mean FR correlation (average weighted degree) per unit across conditions. (**I**) Weighted clustering coefficient per unit across conditions. (**J**) Average shortest path length (largest connected component). (**K**) Normalized betweenness centrality change relative to baseline (percentile bands as in **F**). (**L**) Louvain modularity. (**M**) Weighted rich-club coefficient (φ^w^) as a function of node strength percentile per condition.

### Burst-aligned temporal structure

The SpikeSliceStack provides the foundation for analyzing within-burst temporal dynamics. Population bursts were detected per condition, and activity was aligned to burst peaks using‘align_to_events()’, producing a SpikeSliceStack where each slice captures a single burst window (-250 to +500 ms relative to burst peak). Units were classified into backbone (active in every burst in at least one condition) and non-rigid groups based on burst participation (**Fig. 6A**).^29^ Burst-aligned single unit firing (**Fig. 6B**) and average firing rate heatmaps, ordered by each condition’s median peak time, reveal how the temporal sequence of unit activation changes with dose (**Fig. 6C**).

**Figure 6:**
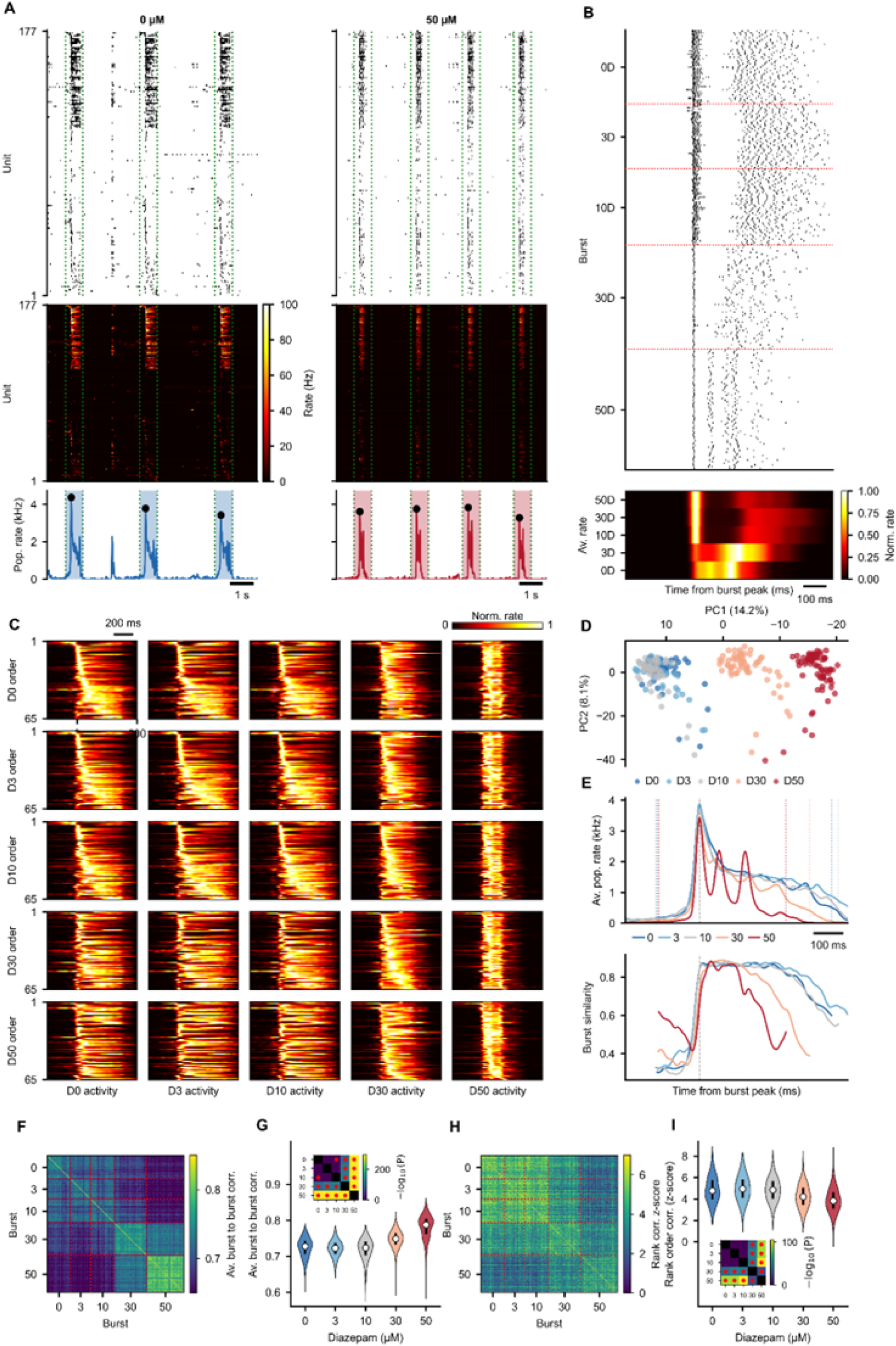
Diazepam disrupts burst-level temporal structure. (**A)** Representative spike rasters, firing rate heatmaps, and population rate traces at 0 µM (left) and 50 µM (right) with detected burst windows highlighted. Units are divided into two groups based on burst participation: Backbone (units firing ≥2 spikes in 100% of bursts in at least one condition) and Non-Rigid (remaining units). Within each group, units are ordered by median firing rate peak time relative to burst peak at baseline. Markers indicate burst peaks and dotted green lines indicate slice boundaries at-250ms to 500ms relative to burst peaks. **(B**) Burst-aligned single-unit spike raster (top) and row-normalized average firing rate heatmap (bottom) for an example unit across all bursts, grouped by condition. Red dashed lines: condition boundaries. (**C**) 5 × 5 grid of burst-aligned average firing rate heatmaps for backbone units. Rows: unit ordering derived from each condition’s median peak time. Columns: firing rate data from each condition. Each unit row-normalized to [0, 1]. (**D**) PCA of burst-to-burst pairwise unit correlation matrices. Each point represents one burst, colored by condition. (**E**) top Burst-aligned average population rate per condition with vertical markers indicating the most conservative burst edges. bottom Burst-to-burst similarity (average slice-to-slice time correlation) per condition, clipped to each condition’s burst edge window. (**F**) Burst-to-burst unit correlation heatmap (mean across units). Red dashed lines separate conditions. (**G**) Within-condition burst-to-burst correlation distributions. Inset: pairwise t-test p-value matrix (−log_10_(*P*); Bonferroni-corrected; red dots: *P* < 0.05). (**H**) Rank-order correlation heatmap (z-scored against shuffled order null-distribution). (I) Within-condition rank-order correlation distributions. Inset: pairwise t-test p-value matrix as in (**H**).

Because each slice is a full SpikeData object, pairwise unit correlations could be computed per burst, yielding a PairwiseCompMatrixStack across bursts. PCA on these burst-level correlation matrices further visualizes how within-burst structure varies across conditions (**Fig. 6D**).

Burst-to-burst similarity was quantified through slice-to-slice time correlations^29^, using (**Fig. 6E**), slice-to-slice unit correlations (**Fig. 6F-G**) and rank-order correlations z-scored against shuffled null distributions (**Fig. 6H-I**), with within-condition distributions tested for cross-condition differences (**Fig. 6G**, **I** insets). Notably, although overall slice-to-slice unit correlations increased with diazepam, specific sequential ordering became less consistent, possibly due to the multi-peak nature of high diazepam concentration bursting (**Fig. 6E**). The analyses in this figure rely on‘align_to_events()’,‘unit_to_unit_correlation()’,‘get_slice_to_slice_unit_corr_from_stack()’,‘get_slice_to_slice_time_corr_from_stack()’,‘get_unit_timing_per_slice()’, and ‘rank_order_correlation()’, highlighting the composability of the SliceStack, where event alignment, per-slice analysis, and cross-slice comparison are all performed through the same interface.

### Latent state dynamics

To move beyond pairwise structure to population-level dynamics, we applied a Gaussian Process Latent Variable Model (GPLVM) to the burst-period firing rates concatenated across all five conditions^40^. The GPLVM assigns each time bin a posterior probability over latent states, capturing the population’s dynamical repertoire without requiring a fixed number of discrete states.

Burst-stitched rasters with GPLVM state overlays show how state occupancy and transitions change between baseline and high-dose conditions (**Fig. 7A**). State-to-state transition matrices, computed only within bursts, reveal dose-dependent changes in dynamical flow (**Fig. 7B**), most notably as a loss of a preserved sequential state ordering upon higher diazepam concentrations, consistent with the rank order statistics reported previously (**Fig. 6H-I**). This loss of temporal state persistence is further denoted by the decrease in average *P*(continuous) traces aligned to burst peaks (**Fig. 7C**). State occupancy distributions (**Fig. 7D**) and per-burst state entropy (**Fig. 7E**) quantify the diversity of visited states and reflect a clear shift in dominant states induced by diazepam. PCA on the instantaneous firing rates provides a complementary low-dimensional view, with burst trajectories colored by GPLVM state (**Fig. 7G-H**) and by condition (**Fig. 7I**), connecting the latent state assignments to the population firing rate manifold. This analysis uses ‘align_to_events()’,‘fit_gplvm()’, and ‘get_manifold()’ to demonstrate how RateData-derived lower dimensional representations and latent state space models can be combined within the same workspace to characterize population dynamics at multiple levels of abstraction.

**Figure 7:**
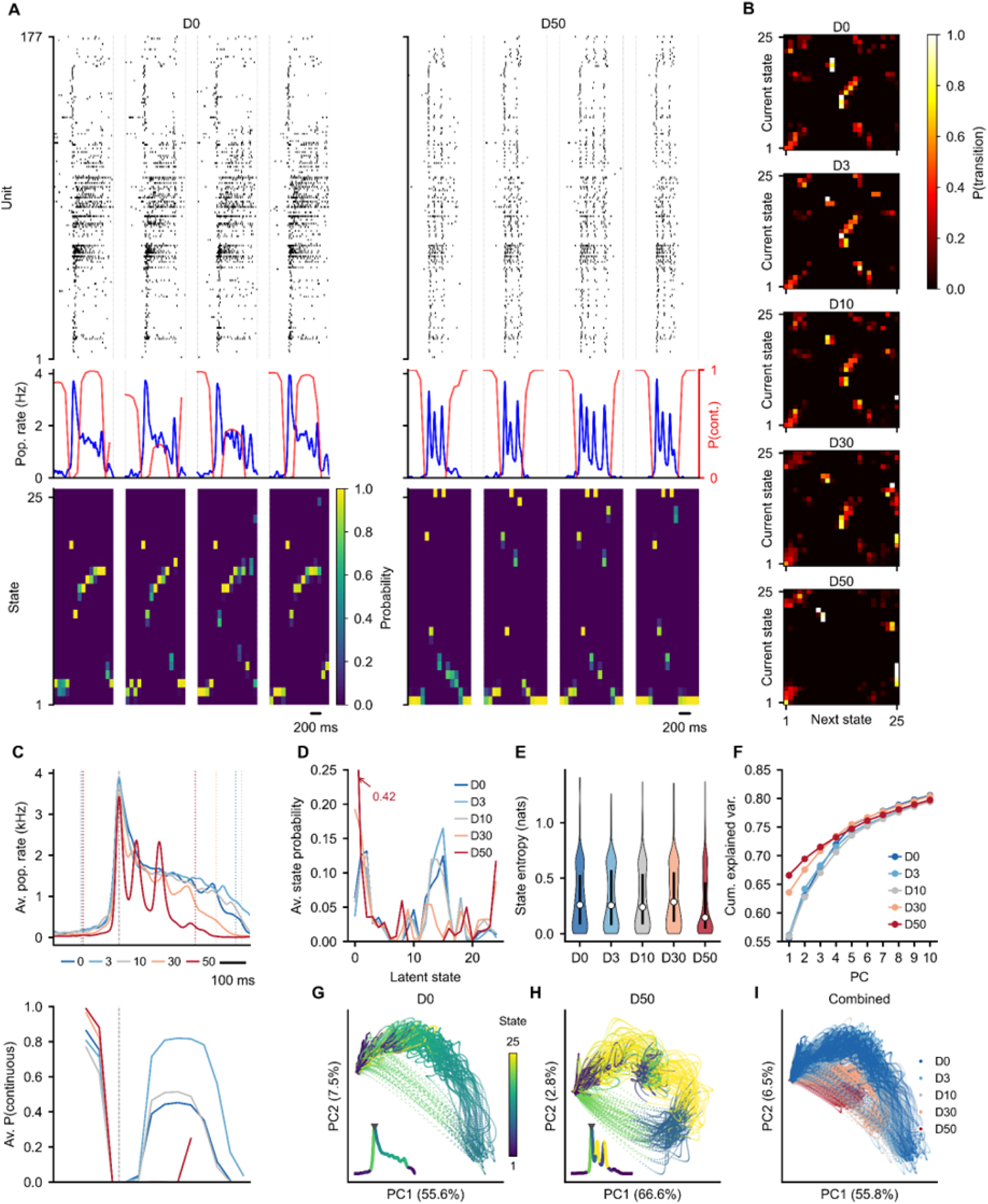
GPLVM latent state analysis reveals dose-dependent changes in burst dynamics. (**A**) top Burst-stitched spike rasters, middle population rate (blue) with P(continuous) overlay (red), bottom GPLVM state probabilities; for 0 µM (left) and 50 µM diazepam (right). Units are ordered based on tuning curve similarity. Bursts are concatenated with silence gaps in between. (**B**) State-to-state transition probability matrices per condition. Only within-burst transitions are counted (cross-burst boundaries excluded). (**C**) top Burst-aligned average population rate (top) and bottom average *P*(continuous) per diazepam concentration. Dotted vertical lines: most conservative burst edges. P(continuous) traces clipped to each condition’s burst edge window. (**D**) Average GPLVM state occupancy probability per condition. (**E**) Per-burst state entropy distributions across conditions. White dots: medians; thick lines: interquartile range. (**F**) Cumulative explained variance from PCA on firing rate data per condition. (**G**, **H**) PC1 vs PC2 of firing rates at 0 µM (**G**) and 50 µM (**H**), with burst time points colored by GPLVM most-likely state. Inset: burst-aligned average population rate colored by state. Shared colorbar: state index. (**I**) Combined PCA embedding across all conditions. Burst time points colored by condition.

The modular data class architecture is designed so that novel analysis methods can be added to the existing framework with minimal effort, inheriting the same composability, workspace persistence, and agentic accessibility. As a result, the library offerings are anticipated to grow over time. The full and up-to-date library contents, including API documentation, are available on GitHub (https://github.com/braingeneers/SpikeLab) and the documentation site (spikelab.braingeneers.gi.ucsc.edu).

## Discussion

The results presented here add to a growing body of evidence that pairing LLMs with domain-specific tools outperforms unassisted models^8,9,13^, demonstrating that this principle extends to multi-step neuroscience analysis pipelines where silent methodological errors are a primary risk. The benchmark results align with previous works reporting that unassisted models fail systematically rather than stochastically, where issues like fabricated API calls, arbitrarily chosen parameters, and implicit simplifications recur across runs.^13,14,15,16,18,19,20^ Many of these errors execute silently without raising exceptions, so repeated runs can reinforce rather than surface the underlying mistake. These are not limitations of reasoning ability; both Opus and Sonnet demonstrated strong capabilities in API exploration, code generation, and debugging. The limitation is the absence of stable, validated building blocks that encode domain expertise. SpikeLab addresses this by providing such building blocks and constraining the model to use them.

### Bounded autonomy as a design pattern

A central finding is that simple text-based constraints towards mandatory library usage, enforcing correctness over efficiency and encouraging clarification-seeking behaviour are able to convert systematic failure modes into reliable behavior. The analysis-implementer skill’s “never assume, ask if unsure” rule converted zero clarification requests across six unassisted runs into a 100% clarification request rate across three SpikeLab runs on an ambiguous task. The “correctness over efficiency” rule prevented silent temporal downsampling that both unassisted models applied without reporting. The “use library methods” rule prevented *ad hoc* method invention that produced three incomparable coupling metrics across three Opus runs (**Fig. 2**).

These constraints do not limit capability, they redirect it. The model retains its flexibility for composing analyses, interpreting results, adapting to new datasets, and generating visualizations. What changes is that the solution space is bounded by validated methods rather than whatever approach the model’s training data happens to favor. This pattern of constraining where correctness matters while preserving flexibility where the model adds value generalizes beyond neuroscience to other domains where LLMs are applied to scientific analysis.^8,9,10^

### The library as persistent expert-validated memory

Without a domain-specific library, every LLM session starts from scratch, forcing the model to reconstruct analytical methods from its training data, with no guarantee that the resulting implementation follows established methodology or matches previous sessions. SpikeLab’s repository addresses this by functioning as a persistent, expert-validated knowledge base.

Unlike prompt-based instructions or in-context examples, library methods are tested, documented, and versioned. They encode domain expertise as durable, reusable code rather than ephemeral context.

The composable data class hierarchy (**Fig. 1**) provides a stable foundation for extending the library. New analytical methods can be added to existing data classes without altering the underlying representations, and they immediately inherit the same slicing, subsetting, workspace persistence, and agentic accessibility as all existing methods. The developer skill closes a feedback loop that makes this extension continuous. Novel analyses conducted through the analysis-implementer can be integrated back into the library, where they undergo testing, review, and documentation before becoming available to all users and agents. This enables community contributions through the same agentic workflow. Researchers add new methods that are immediately accessible, without requiring downstream users to understand or reimplement the underlying computations. Over time, the library is positioned to grow as a shared repository of validated analytical procedures that persists across sessions, users, and model versions, following the model of community-driven analysis ecosystems that have emerged in other fields.^41,42^

To preserve this guarantee across releases, the library commits to backwards-compatible evolution where possible: existing data class signatures, method names, and return shapes are maintained across versions, so analyses and skill-generated code from prior sessions can continue to run unchanged.

### Workspace as session memory

Beyond the library itself, the AnalysisWorkspace provides a form of session-level memory that complements the library’s role as a persistent knowledge base. Each workspace stores intermediate results, cached computations, and multiple recordings in a single HDF5 file, alongside analysis scripts and an analysis log that documents the methods applied, findings, and open questions. Together, the analytical architecture presented here forms a self-contained record of an analysis session that can be shared with collaborators or resumed by a new agent without loss of context.

Our presented framework also supports collaborative and parallel workflows. Multiple recordings can be analyzed within a single workspace, enabling cross-recording comparisons without format conversion. The workspace merge operation allows results from independent agent sessions to be combined. For instance, two agents could implement different analytical workflows in parallel and their workspaces can be merged afterward. This positions the workspace not just as a caching layer but as an infrastructure for scalable, multi-agent analysis workflows.

### Cost-effectiveness

The benchmark also revealed a practical implication for resource allocation. Opus, the most capable model tested, was the most expensive condition (An estimated $1.95 per session on average at high reasoning effort) yet did not outperform SpikeLab-assisted Sonnet (An estimated $0.69 per session on average at medium effort) on the scorecard. Investing in domain-specific tooling for a less expensive model yielded better results than deploying a more expensive model from first principles. As model costs continue to decline^43^, the relative value of domain-specific structure over raw model capability is likely to increase.^13,44^

### Limitations

Several limitations should be considered when interpreting these results. The benchmark evaluated a single dataset (IBL auditory cortex) with four tasks and three runs per condition. While a broader task set and additional datasets would strengthen the generalizability of the claims, the same classes of errors, including *ad hoc* method invention, silent simplification and unreported decisions, resurfaced independently across different tasks and runs. This suggests that these are systematic tendencies of unassisted models rather than artifacts of the specific benchmark. The benchmark also evaluated a single model family (Claude); whether the same patterns hold for other model families remains untested, though studies benchmarking multiple model families have observed similar failure modes across architectures^13,14,15,16^, suggesting the underlying tendencies are general rather than model-specific.

SpikeLab’s current scope is limited to spike train data and does not extend to other recording modalities such as calcium imaging, EEG, or fMRI. With the framework of tool-based encoding of scientific domain expertise validated, adding novel tools for different data types is anticipated to be relatively straightforward. Already in the current version, the library supports a broad range of analyses spanning single-unit statistics, pairwise correlation and network structure, and population-level dynamics (**Fig 4-7**). Moreover, as demonstrated in **Fig. 3**, these analyses can be applied to diverse data sources including *in vivo* recordings from different species, different recording technologies, and *in vitro* preparations without modification. The architectural pattern of composable data structures, bounded agentic skills and workspace persistence is not inherently modality-specific and could serve as a template for similar frameworks in other domains.

### Future directions

Several extensions are natural. Expanding the library to support additional data modalities such as calcium imaging, local field potentials and multi-modal recordings would broaden applicability while preserving the same architectural guarantees. SpikeLab already integrates with neuroscience data standards through NWB data loaders^45^ and a dedicated IBL data bridge^33^, which can also serve as a source of diverse, high-quality recordings for developing and testing analysis pipelines before acquiring new data. The skill system could be extended with additional specialized roles like a literature-aware skill that grounds analytical choices in published methods or a statistical advisor that can recommend appropriate tests and help shape experimental design.^46^

More broadly, the text-to-analysis paradigm demonstrated here suggests a model for AI-assisted science in which domain expertise is encoded not in the model’s weights but in the tools it is given access to. As LLMs become more capable, the value of such tools shifts from compensating for model limitations to ensuring that increased capability is channeled through validated methodology. The goal is not to replace the researcher but to provide a structured environment in which both human expertise and model capability can be applied reliably.

## Methods

### Library implementation

SpikeLab is implemented in Python and installable via‘pip install spikelab’. The library is structured around five core data classes: SpikeData, RateData, PairwiseCompMatrix, PairwiseCompMatrixStack, and RateSliceStack/SpikeSliceStack. These classes form a composable hierarchy for representing spike trains, firing rates, pairwise comparisons, and event-aligned trial stacks. A workspace persistence layer (AnalysisWorkspace) provides HDF5-based caching of intermediate results addressed by (namespace, key) pairs. Data loaders support HDF5, NWB, KiloSort/Phy, SpikeInterface, and pickle formats, as well as the International Brain Laboratory (IBL) public dataset via the ONE API. Exporters write SpikeData to HDF5, NWB, CSV, and KiloSort formats for integration with external pipelines.

PairwiseCompMatrix objects can be converted to weighted NetworkX graphs for network analysis. All spike times are stored and processed in milliseconds throughout the library. Source code is available at https://github.com/braingeneers/SpikeLab.

The library has no mandatory dependencies beyond NumPy, SciPy, and Matplotlib. Optional dependencies are imported at call time and raise informative errors if absent: h5py for workspace persistence, boto3 for S3 access, numba for accelerated computations, umap-learn for UMAP dimensionality reduction, and poor_man_gplvm^40^ with jax for Gaussian Process Latent Variable Models.

Performance-critical routines like spike time tiling coefficient (STTC), pairwise nearest-spike latencies, and spike-triggered population rate have optional Numba-accelerated kernels that are used automatically when Numba is installed. The degree-preserving raster shuffle (‘randomize()’)^27^ operates on a binary raster matrix at user-specified bin resolution (default 1 ms) and preserves both per-unit spike counts and per-timebin population rates via iterative swap operations.

### Agentic skill system

The skill system consists of markdown instruction files that are loaded into the LLM’s context when invoked. Skills are not executable code but rather structured natural language instructions that shape the model’s behavior within a conversation session. Four skills are shipped with the library (in‘.agent/skills/’):

- The analysis-implementer skill defines file scope boundaries, mandates library method usage over custom implementations, enforces correctness-over-efficiency priorities, requires clarification on ambiguous scientific choices, and prescribes workspace caching and analysis logging practices.
- The educator skill provides read-only access for concept explanation and result interpretation, without the ability to create, modify, or execute any files.
- The spike-sorter skill manages the upstream spike sorting pipeline, wrapping SpikeInterface^24^ and external sorting algorithms (e.g., Kilosort^25^) with automated curation and provenance tracking. It outputs curated SpikeData objects ready for analysis.
- The developer skill governs library development like adding new methods, modifying existing code, and integrating user-provided analysis routines into the library. It operates under strict scope boundaries (library source only, no tests or analysis scripts) and requires reading the repo maps before making changes.

A separate orchestrator skill spikelab lives outside the repository and serves as the user-facing entry point. It handles cloning the repository, installing the library, generating the repo maps, and routing tasks to the appropriate in-repo skill. This skill is provided alongside the manuscript as a standalone file that can be added to any compatible agentic coding environment.

All skills reference repo map files, two markdown documents (REPO_MAP.md and REPO_MAP_DETAILED.md) that provide a condensed quick reference and a full API reference, respectively, for the entire library. These are maintained alongside the codebase and document every class, method, signature, parameter, and return type. Because the full repository is too large to load into context, the repo maps serve as the agent’s primary orientation mechanism.

The agent consults them to locate relevant classes or methods, then greps the underlying source files to read only the specific sections it needs before writing a custom analysis script. This treats the codebase as an external environment the agent queries on demand rather than as context to be ingested wholesale, drawing on the framing of Recursive Language Models which similarly handle oversized inputs by storing them externally and having the model programmatically inspect relevant fragments.^14^ A practical consequence is that the repository can continue to grow without the context window becoming a bottleneck. Adding new modules or methods expands what the agent can do but does not increase what must be loaded into context for any single task. Beyond scalability, this pattern also improves accuracy. By consulting the repo maps before writing code, the agent verifies method signatures and parameters, contributing to SpikeLab’s reduced error rate compared to models operating without library documentation.

The MCP server exposes library methods as stateless tools accessible via the Model Context Protocol^36^, providing an alternative programmatic interface that does not require a skill-based agentic session. Any MCP-compatible client, including coding agents, IDE extensions, or custom pipelines, can call these tools directly. Each tool reads from and writes to the workspace via‘(workspace_id, namespace, key)’ addressing.

### Benchmark protocol

#### Conditions

Three conditions were compared on a standardized four-task analysis pipeline:

1. Opus: Claude Opus 4.6 at high reasoning effort, bare conda environment (Python 3.10), no pre-installed scientific packages
2. Sonnet: Claude Sonnet 4.6 at medium reasoning effort, same bare environment
3. SpikeLab: Claude Sonnet 4.6 at medium reasoning effort, same bare environment plus pre-installed SpikeLab with all dependencies

All conditions used Claude Code (v2.1.81) as the agentic coding interface. Each condition was run three times on consecutive days (run dates: 2026-03-24/25/26), yielding 9 sessions total. All conditions received identical task prompts (Fig. 2A), with the sole exception that the SpikeLab condition was prefixed with the skill invocation command on Task 1. Opus and Sonnet were responsible for installing their own packages (numpy, scipy, matplotlib, etc.) from the bare environment.

#### Tasks

The four tasks represent a progressive neuroscience analysis workflow on publicly available data from the International Brain Laboratory (IBL). Verbatim prompts:

1. Task 1: “Download the IBL dataset with the most good units in auditory cortex and plot a rasterplot of activity in minute 20 to 21, together with the population rate and instantaneous firing rates. Order units from highest to lowest average firing rate over the whole recording.”
2. Task 2: “Compute the spike time tiling coefficient (STTC) between all pairs of neurons (delta = 20ms). Apply z-score normalization using 10 shuffled datasets to account for possible effects of firing rates. Run this analysis separately on the cue and no-cue parts of the recording. Plot: (1) the z-scored STTC pairwise matrix as a heatmap for each condition, ordered by average correlation strength, (2) a scatterplot comparing cue vs no-cue z-scored STTC values for each neuron pair, with marginal histograms on both axes.”
3. Task 3: “Compute the spike-triggered population rate for each unit and extract the zero-lag coupling strength. Select the unit closest to the median population coupling strength. For that unit, create an event-aligned spike raster plot across all stimulus trials (200ms pre-stimulus to 500ms post-stimulus). Below the raster, plot the average population firing rate across all trials as a PSTH with a shaded SEM band.”
4. Task 4: “Compute instantaneous firing rates and perform PCA on the population rate matrix. Fit a latent state space model to the spike data. Plot the first two principal components as a trajectory, coloring each point by the most likely state from the fitted model. Report the variance explained by the first 5 components.”

Task 2 was deliberately underspecified. The phrase “run this analysis separately on the cue and no-cue parts of the recording” admits multiple valid interpretations of “no-cue”: inter-trial intervals (animal in task context but no trial active), catch trials (zero-contrast trials with full task engagement), or pre/post-task periods (no task demands), to test whether models would seek clarification or proceed with an unexamined assumption.

#### Evaluation

Each run was evaluated against a 14-issue scorecard (**Fig. 2D**) spanning data discovery (3 issues), methodological correctness (5 issues), and scientific documentation practices (3 issues), plus 3 task-specific model selection issues. Each issue was scored per run as correct (green), partially correct (orange), or incorrect/absent (red). Scorecard issue descriptions and supporting code examples are provided in Supplementary Materials. Raw session transcripts and all generated analysis scripts and figures are available in Supplementary Data.

#### Tool call quantification

Tool calls were extracted from Claude Code session transcripts and classified into seven categories: failed executions (non-zero exit code), inline exploration (python-c one-liners), script writes (Write/Update tool calls), successful script runs, environment setup (pip install, mkdir, skill loading), file reads (Read tool calls to output files and figures), and documentation reads (Search/Grep/Read calls to SpikeLab library source code and repo maps).

Several classification rules were established to resolve ambiguous cases consistently across all transcripts:

- Counting granularity: Each tool invocation was counted as one action regardless of internal fan-out. In particular, an Explore agent call, a subagent that may internally perform multiple grep and file read operations, was counted as a single documentation read, and a Read call accessing multiple files was counted as a single file read. This ensures consistent treatment across tool types: counts reflect strategic decisions, not the number of operations each tool performs internally. Consecutive Update calls to the same file with no intervening reasoning were counted as a single script write.

- Error handling: Any tool call that exited with a non-zero exit code was classified as a failed execution, regardless of whether it was a‘python-c’ one-liner, a full script run, or a compound command. When a model killed a stuck background process, the kill command was classified as a failed execution and the original background launch was excluded to avoid double-counting.

Tool calls that produced no effect (e.g., a failed Edit where the file was unchanged) were excluded.

- Monitoring exclusions: Background monitoring polls like‘sleep N && cat/tail’ commands checking whether a running script had finished, and‘ps’/‘tasklist’/‘wmic’process checks were excluded entirely, regardless of whether they succeeded or errored, as they reflect the runtime environment rather than the model’s analytical strategy. Background task completion notifications for already-counted launches were not counted separately.

- Environment and setup: Package import checks (‘python-c “import spikelab; print(’OK’)”’) and Task Output/Stop Task commands were classified as environment setup.

- Write classification: Writes to‘ANALYSIS_LOG.md’ were counted as script writes.

All 252 values (9 sessions x 4 tasks x 7 categories) were verified through independent agentic re-counting of every transcript, with each count traced to specific transcript line numbers. Raw session transcripts and line-by-line evidence are available in Supplementary Data.

### Analysis methods

#### Burst detection:‘get_bursts()’ (Figs. 3–7)

Two smoothed population rate traces were computed from the binned population spike count (1 ms bins): a coarse trace (boxcar width 20 ms, Gaussian sigma 100 ms) for peak detection and edge finding, and a high-resolution trace (boxcar width 8 ms, Gaussian sigma 8 ms) for refining peak locations within detected burst windows. Burst peaks were identified by thresholding the coarse trace at a species-specific RMS multiplier: 1.25x for mouse, 1.5x for human, 2.5x for organoid. Minimum inter-burst interval was set to 1000 ms for all. Burst edges were defined using peak-to-trough baseline estimation: the edge threshold was set at 0.2x the difference between the peak amplitude and the higher of the two flanking troughs on the coarse trace.

Final burst peak times were refined to the maximum of the high-resolution trace within the detected edge window.

#### Spike time tiling coefficient:‘spike_time_tilings()’ (Figs. 2, 5)

STTC was computed for all unit pairs using a 20 ms coincidence window.^30^ Z-score normalization used 10 degree-preserving shuffled datasets generated by‘spike_shuffle_stack()’ following Okun et al.^27^, which preserves both per-unit spike counts and per-time bin population rates.

#### Population coupling:‘compute_spike_trig_pop_rate()’ (Figs. 2, 3, 4)

Spike-triggered population rate (stPR) was computed following Bimbard et al.^38^, building on Okun et al.^39^ For each unit, the leave-one-out population rate was computed, baseline-subtracted, normalized by the sum of mean firing rates of all other units, low-pass filtered at 20 Hz (Butterworth), and evaluated over a +/- 80 ms lag curve. Zero-lag and maximum-lag coupling strengths were extracted per unit.

#### Instantaneous firing rates:‘resampled_isi()’ (Figs. 3–7)

Instantaneous firing rates were estimated using ISI-interpolated rates at 1 ms resolution with Gaussian smoothing (10ms sigma). This method estimates rate from the reciprocal inter-spike interval interpolated at arbitrary time points, adapting naturally to local firing patterns and avoiding discretization artifacts.

#### Pairwise firing rate correlations:‘get_pairwise_fr_corr()’ (Fig. 5)

Pairwise firing rate correlations were computed from instantaneous firing rate traces using cross-correlation with a maximum lag of 350 ms, following van der Molen et al.^29^ Correlation matrices were stored as PairwiseCompMatrix objects supporting network graph conversion, thresholding, and spatial visualization.

#### Network graph metrics:‘to_networkx()’ (Fig. 5)

Functional connectivity graphs were constructed from pairwise FR correlation matrices using NetworkX. Metrics computed: mean weighted degree (node strength), weighted clustering coefficient, characteristic path length (largest connected component), betweenness centrality, Louvain modularity, and weighted rich-club coefficient as a function of node strength percentile.

#### Burst-aligned analysis:‘align_to_events()’,‘get_slice_to_slice_unit_corr_from_stack()’,‘get_slice_to_slice_time_corr_from_stack()’,‘unit_to_unit_correlation()’,‘rank_order_correlation()#x2019; (Figs. 6, 7)

Burst peak-aligned RateSliceStacks and SpikeSliceStacks were created by aligning to burst peak times with-250 ms pre and +500 ms post windows. Burst-to-burst unit correlations, slice-to-slice time correlations, unit-to-unit correlation PCA (2 components), and rank-order correlations (z-scored against 100 shuffled permutations) were computed on the combined slice stacks.

Units were classified into two groups: Backbone (units firing >= 2 spikes in 100% of bursts in at least one condition) and Non-Rigid (all remaining units). Within each group, units were ordered by median firing rate peak time relative to burst peak at baseline.

#### GPLVM:‘fit_gplvm()’ (Fig. 7)

A Gaussian Process Latent Variable Model^40^ was fitted on burst-period firing rates concatenated across all five diazepam conditions with 100 ms silence gaps between bursts. The model used 25 latent bins with 50 ms bin size. Per-condition results were extracted by slicing the combined model output at condition boundaries. State entropy was computed as the Shannon entropy of the GPLVM state posterior per burst. State-to-state transition probability matrices excluded cross-burst boundaries.

#### PCA:‘get_manifold()’ (Figs. 6, 7)

Principal component analysis was performed on instantaneous firing rate matrices (ISI-interpolated, Gaussian-smoothed) using 2 components. Per-recording and combined (all conditions concatenated) embeddings were computed. Cumulative variance explained was reported for the first 5 components.

#### Sliding-window analysis:‘frames()’ (Fig. 3)

Population coupling was computed across sliding 2-minute windows (120 s window, 10 s step = 110 s overlap) over the human recording. Per-window coupling was computed via spike-triggered population rate at zero lag. Units were included in a window if they fired >= 12 spikes. Normalized change relative to the first window was computed using symmetric normalization: N = (x’ - x_0_’) / (x’ + x_0_’), where x’ = max(x, 0).

#### Burst sensitivity analysis”‘burst_sensitivity()’ (Figs. 3, 4)

Burst detection threshold sensitivity was assessed by sweeping the RMS multiplier from 1.0 to 5.0 (step 0.1 for **Fig. 3**; step 0.25 for **Fig. 4**) while holding minimum inter-burst interval (1000 ms) and edge threshold (0.2x) constant. Burst counts per minute were reported as a function of threshold.

#### Data Availability: IBL auditory cortex (benchmark, Fig. 2; cross-data-type, Fig. 3)

Mouse auditory cortex data from the International Brain Laboratory Brain-Wide Map.^33^ Session‘7f5df7eb’, probe insertion‘0c15a331’: 208 good-quality units in auditory cortex subregions AUDv and AUDd. Recording duration: 77.6 minutes. The dataset is publicly available via the IBL ONE API (https://docs.internationalbrainlab.org/notebooks_external/one_quickstart.html).

**Human anterior temporal lobe (Fig. 3).** 152 units recorded in vivo from human anterior temporal lobe (participant NIH062) with dual 64-channel Utah arrays during a visual stimulus paradigm.^34^ Recording split into cue (stimulus-driven, 935 s) and no-cue (spontaneous, 2318 s) periods at 5 s after the last stimulus offset. **Brain organoids (Figs. 3–7).** Human forebrain organoid recorded on a multi-electrode array (MEA). Diazepam dose-response paradigm: 5 concentrations (0, 3, 10, 30, 50 uM), each recorded in a separate ∼180 s session. 177 units matched across all conditions. Spike sorting was performed using the in-repo Kilosort2 pipeline with default parameters; all five recordings were concatenated before sorting to maintain consistent unit identities across conditions. Original recordings from Sharf et al.^35^; raw and spike sorted data can be found here https://datadryad.org/dataset/doi:10.25349/D9031Z.

## Code Availability

All analyses were performed using SpikeLab v0.1.0 with Python 3.10. Figure generation code, analysis scripts, computed workspaces, and raw benchmark transcripts are available in the manuscript repository at https://github.com/braingeneers/SpikeLab-manuscript. The complete figure reproduction pipeline can be executed from ‘manuscript/figure_code/’ using ‘python-m fig{N}.assemblè. The distributable SpikeLab skill that handles installation, environment setup, and delegation to specialized analysis, sorting, education, and development roles is included in Supplementary Data and can be downloaded at 10.5281/zenodo.19714691.

## Supporting information

Spikelab skill

## Acknowledgments

This study was supported by the National Science Foundation (NSF) Emerging Frontiers in Research and Innovation under award (NSF 2515389 to T.S., M.T., and D.H.), the Brain and Behavior Research Foundation Young Investigator Award (to T.S.), the Schmidt Futures Foundation SF857 and the National Human Genome Research Institute under Award number RM1HG011543 (D.H. and M.T.), the National Institute of Mental Health of the National Institutes of Health under Award Number 1U24MH132628 (D.H.), the National Science Foundation under award number NSF 2034037 (M.T), and NSF 2134955 (M.T. and D.H). This work was also supported in part by National Science Foundation (NSF) awards CNS-1730158, ACI-1540112, ACI-1541349, OAC-1826967, OAC-2112167, CNS-2100237, CNS-2120019, the University of California Office of the President under Award number M25PR9045, and the University of California San Diego’s California Institute for Telecommunications and Information Technology/Qualcomm Institute.

## Author contributions

Conceptualization: T.v.d.M., T.S. and D.H., Methodology: T.v.d.M., A.S., A.R. and D.P., Software: T.v.d.M., L.C., K.H., O.B., M.L. and J.G., Validation: T.v.d.M., Formal analysis: T.v.d.M., Resources: T.S., D.H., M.T. and K.K., Writing – original draft: T.v.d.M., Writing – review & editing: T.S., Visualization: T.v.d.M., Supervision: T.v.d.M. and T.S., Funding acquisition: T.S., D.H., M.T. and K.K. All co-authors provided their feedback on the final version of the manuscript

## Supplementary materials

### Spikesorting module

SpikeLab includes a spike sorting module (‘spikelab.spike_sorting’) that provides a text-to-analysis interface for the spike sorting stage of the electrophysiology pipeline. Built on top of SpikeInterface^1^, the module supports three sorting backends, Kilosort2^2^, Kilosort4^3^, and RT-Sort^4^, and implements a multi-stage curation workflow that can filter units based on firing rate, ISI violation ratio, signal-to-noise ratio, spike count, and waveform spatial consistency. A dedicated spike-sorter skill wraps this module for agentic access, extending the text-to-analysis framework from the analysis stage back to the data preprocessing stage.

### Sorting backends

The module implements a backend abstraction layer (‘SorterBackend’) that defines three operations: load and preprocess a recording, run the sorter, and extract per-unit waveforms. Each backend maps these operations to a specific sorting algorithm while exposing a unified interface to the pipeline.

Kilosort2 is a template-matching algorithm that requires MATLAB. SpikeLab supports two execution modes: a local installation with a user-provided Kilosort2 source path, and a Docker container that bundles the compiled MATLAB Runtime (no MATLAB license required). The Docker image is selected automatically based on the host GPU’s NVIDIA driver version.

Kilosort4 is a pure-Python implementation using PyTorch. It runs locally with a CUDA-enabled PyTorch installation or in a Docker container. Kilosort4 uses updated template-matching and clustering algorithms compared to Kilosort2 and requires no MATLAB dependency.

RT-Sort is a deep-learning-based sorter that detects spikes using a trained neural network model and clusters them based on propagation sequence similarity across the electrode array. RT-Sort requires PyTorch with CUDA and is not available through Docker. The trained RT-Sort object is serialized to disk after sorting, enabling reuse in stimulation-aware sorting (see below).

All three backends produce SpikeInterface-compatible sorting objects, which are then processed through the same downstream pipeline: waveform extraction, SpikeData construction, and curation. The backend architecture is designed for extensibility. Adding a new sorting algorithm requires only implementing the sort step itself; everything else (recording loading, configuration management, format detection, folder setup, logging, waveform extraction, SpikeData conversion, curation, figure generation, compilation, and pickle serialization) is handled by the shared pipeline and applies automatically to any registered backend.

### Recording handling

The module accepts recordings in Maxwell HDF5 (‘.raw.h5’) and NWB (‘.nwb’) formats natively. In addition, any pre-loaded SpikeInterface‘BaseRecording’object can be passed directly as input in place of a file path.^1^ Because SpikeInterface provides recording extractors for a wide range of electrophysiology file formats, including Open Ephys, Intan, Neuralynx, Plexon, Blackrock, and others, this indirectly extends the module’s format support to the full SpikeInterface ecosystem.

Two entry points handle different recording scenarios.‘sort_recording()’accepts a list of recording file paths and sorts each independently. When a directory is passed instead of a file, all recordings in the directory are concatenated into a single continuous trace (using alphabetic ordering of file names), sorted together, and split back into per-file SpikeData objects with epoch-specific waveform templates. ‘sort_multistream()’handles multi-well Maxwell recordings by iterating over user-specified well identifiers (stream IDs) and sorting each well independently. Similarly, passing a directory of multi-well recordings will concatenate the different recordings before sorting, at a well by well level.

Both entry points support time-windowed sorting: a single contiguous window (‘start_time_s’, ‘end_time_s’), the first N minutes of a recording (‘first_n_mins’), or multiple disjoint windows (‘rec_chunks_s’) that are concatenated before sorting and split afterward.

### Curation

Curation is optional and can be disabled entirely, applied during sorting with configurable thresholds, or performed post-hoc on existing SpikeData objects. The pipeline supports five curation criteria: minimum firing rate (default: 0.05 Hz), maximum ISI violation ratio (default: 1.0%, computed at a 1.5 ms threshold), minimum signal-to-noise ratio (default: 5.0), minimum spike count (default: 50), and maximum normalized waveform standard deviation (default: 1.0), which removes units with spatially inconsistent waveform shapes. All thresholds are configurable through a ‘CurationConfig’dataclass. For concatenated recordings, curation can be restricted to a single epoch, allowing quality assessment on the most representative segment while retaining spikes across the full recording.

When curation is applied, the pipeline produces a serializable curation history that records: the initial unit set, each criterion with its threshold and per-unit pass/fail outcome, per-unit quality metrics, and the final curated unit set. This history is saved as JSON alongside the results.

Post-hoc curation methods on SpikeData objects allow researchers to apply additional or alternative filters after sorting without rerunning the pipeline. Each method returns a new SpikeData object and per-unit metric arrays, preserving the original data.

### SpikeData construction and provenance

The pipeline converts sorted waveform data into SpikeData objects with full provenance metadata. Each SpikeData object stores spike trains in milliseconds (consistent with the library convention) along with per-unit neuron attributes and recording-level metadata. Recording-level metadata includes the source file path, sorting backend identifier, sampling frequency, electrode positions, and in case of concatenated recordings, epoch boundaries in both samples and milliseconds with the original file names. This metadata is sufficient to trace any SpikeData object back to its source recording and sorting parameters.

Per-unit neuron attributes include: the original cluster ID from the sorter, the maximum-amplitude channel and its electrode coordinates, waveform templates (1D on the max channel, 2D across all channels, and a windowed version), peak amplitude and per-channel amplitudes, signal-to-noise ratio, normalized waveform standard deviation, waveform polarity (positive or negative peak), and the raw spike train in samples for lossless round-tripping. For concatenated recordings, per-epoch average templates are stored separately, enabling drift assessment across recording segments.

The resulting SpikeData objects are directly compatible with all downstream analysis methods described in the main text: firing rate computation, event-aligned slicing, pairwise comparison matrices, population coupling, and latent variable models. This closes the loop between raw recordings and the analysis pipeline.

### Quality control figures

The pipeline optionally generates diagnostic figures at two stages. Before curation, per-unit figures are generated for every detected unit containing an ISI histogram (0–100 ms), a spatial waveform footprint across electrode positions, and a max-channel overlay showing individual spike traces with the mean waveform. After curation, these figures are sorted into‘curated/’and‘failed/’subdirectories, providing a visual audit trail for every curation decision.

Post-curation figures include: a bar plot comparing total versus curated unit counts, a scatter plot of normalized waveform STD versus spike count with curation threshold lines, stacked waveform templates grouped by polarity, four-panel quality metric histograms (SNR, firing rate, spike count, ISI violations. All distributions are computed on the full pre-curation population with threshold lines overlaid), and a raster plot with population rate for the first 30 seconds of the recording.

### Stimulation-aware sorting

For experiments involving electrical stimulation of neural tissue, the module provides a two-step workflow: propagation sequences are first detected in an intrinsic activity recording using RT-Sort^4^ and then applied to sort spikes in the artifact-contaminated stimulation recording.

In Step 1, a baseline recording of intrinsic (unstimulated) activity is sorted using RT-Sort (or a period of the stimulation recording with no stimulations). The resulting RT-Sort object, containing detected propagation sequence templates, is serialized to disk. In Step 2,‘sort_stim_recording()’processes the stimulation recording using the pre-trained sequences. This function first recenters logged stimulation times to actual artifact peaks in the voltage traces (correcting for timing jitter between the stimulation controller and the recording system), then removes stimulation artifacts using per-event polynomial detrending adapted from SALPA [Wagenaar & Potter, J Neurosci Methods 2002]. The polynomial fit captures the slow artifact waveform while preserving fast spike waveforms that co-occur with the artifact. The cleaned traces are then sorted using the pre-detected RT-Sort sequences, and spikes are aligned to the corrected stimulation times.

The output is a SpikeSliceStack aligned to stimulation events, enabling direct analysis of stimulus-evoked responses with the same event-aligned methods used throughout the library. Sequential stimulation protocols (e.g., burst stimulation, paired-pulse) are handled by dynamically extending the blanking region around each stimulation event. Individual pipeline components (artifact time recentering and artifact removal) are also available as standalone functions for custom workflows.

### Spike-sorter skill

The spike-sorter skill wraps the sorting module for agentic access, extending the text-to-analysis framework from the analysis stage (covered by the analysis-implementer skill) back to the data preprocessing stage. The top-level SpikeLab skill automatically routes spike sorting requests to the spike-sorter skill based on the user’s natural language prompt so that users do not need to select the correct skill manually. The skill operates under the same bounded autonomy constraints as the analysis-implementer.

File boundaries: Raw recording files are read-only. The skill never modifies, moves, or deletes source data. Sorting scripts and results are written to a user-confirmed results directory, never to the library source tree.

Analysis boundaries: The skill is restricted to sorting and output quality assessment (unit counts, SNR and firing rate distributions, waveform template inspection, and curation outcomes). For any downstream analysis, the skill directs the user to the analysis-implementer skill with the curated SpikeData pickle as the handoff point.

Clarification-seeking: The skill does not assume recording formats, electrode configurations, sorter choices, or curation thresholds. It asks the user to confirm: the recording format, whether the experiment involves electrical stimulation, single versus multi-well layout, the sorting backend, and any non-default curation parameters.

Mandatory reporting: After every sorting run, the skill generates a markdown sorting report with the curation outcome (raw versus curated unit counts, total spikes, mean firing rate, mean SNR) at the top, followed by pipeline settings, stage-by-stage timing, and unit quality distributions.

This report, combined with the curation history JSON and quality control figures, provides a complete audit trail from raw recording to curated SpikeData.

Downstream handoff: The curated SpikeData pickle (‘sorted_spikedata_curated.pkl’) serves as the bridge between the spike-sorter and analysis-implementer skills. A user can sort recordings in one session and load the results for analysis in a subsequent session, with full provenance metadata preserved. This two-skill design separates spike sorting (where parameter choices depend on the recording hardware and tissue type) from analysis (where method choices depend on the scientific question), while maintaining a single data structure, SpikeData, as the interface between them.

### Batch-jobs module

SpikeLab includes a batch jobs module (‘spikelab.batch_jobs’) that submits analysis and spike sorting workloads to remote Kubernetes clusters. The module is an optional dependency (‘pip install spikelab[batch-jobs]’) and is intended for compute-intensive workflows, like long-running spike sorting, large parameter sweeps or fitting latent-variable models, that exceed the resources of a typical analysis workstation. A natural-language instruction file (‘INSTRUCTIONS.md’) ships with the module, allowing the analysis-implementer and spike-sorter skills to deploy jobs on the user’s behalf without leaving the text-to-analysis workflow.

### Submission modes

Three submission entry points cover the common deployment patterns:

‘submit_workspace_job()’saves an‘AnalysisWorkspacè to disk, bundles it with a user-supplied analysis script, uploads the bundle to S3-compatible storage, and submits a Job that re-loads the workspace, runs the script, and writes the updated workspace back to S3.

‘submit_sorting_job()’bundles a list of recording paths together with a ‘SortingPipelineConfig’(or a preset name) and runs the spike sorting pipeline (Supplementary materials: Spikesorting module) on the cluster.

‘submit_prepared_job()’ submits a Job without bundling artifacts, for users who manage their own input/output paths. All three modes return a ‘SubmitResult’containing the job name, rendered manifest, run identifier, and S3 prefixes for outputs and logs.

The complementary ‘retrieve_result()’method downloads the outputs after completion and reconstructs an‘AnalysisWorkspacè. For sorting jobs, each recording’s curated SpikeData is loaded into a separate workspace namespace, preserving the per-recording structure of the input.

### Cluster profiles

Cluster-specific configuration is decoupled from job specifications through the‘ClusterProfilè model. A profile bundles namespace, default CPU/GPU images, S3 prefix, namespace hooks (credential mounts and environment variables), affinity/tolerations, storage path templates, and policy thresholds. A generic ‘defaults’profile ships with the package, and users can supply custom profiles as YAML files via ‘--profile-filè, allowing the same job spec to be deployed to different clusters by switching profiles only. Namespace hooks automatically mount Kubernetes secrets and inject credential environment variables (e.g., ‘AWS_SHARED_CREDENTIALS_FILÈ,‘KUBECONFIG’), so containers do not need to be root or carry baked-in credentials.

### Policy preflight

Every submission runs through a configurable policy engine (‘evaluate_policy’) before the manifest is rendered. The default rules cap interactive GPU count, cap maximum runtime (default: 14 days), warn when CPU/memory requests and limits diverge, and block batch commands that resemble idle placeholders (‘sleep infinity’, bare ‘sleep’,‘sleep’). Findings are tagged ‘PASS’,‘WARN’, or‘BLOCK’; a ‘BLOCK‘raises‘RuntimeError’unless the caller passes‘allow_policy_risk=Truè. Thresholds are read from the active profile, so different clusters can enforce different rules without code changes.

### Artifact packaging and storage

Bundles are deterministic ZIP archives produced by‘package_analysis_bundle()’, which records SHA-256 hashes for every input file in a manifest alongside the bundle. Bundles, outputs, and logs are addressed through the profile’s ‘StoragePathTemplates’(‘{prefix}/inputs/{run_id}/…’,‘{prefix}/outputs/{run_id}/…’,‘{prefix}/logs/{run_id}/…’), giving each run a self-contained S3 namespace that can be re-downloaded, archived, or shared. The‘S3StorageClient’ works with any S3-compatible endpoint via the profile’s‘endpoint_url’ and‘region_namè, including non-AWS object stores.

### CLI and Python API

The same operations are exposed through both a CLI (‘spikelab-batch-jobs’) and a Python API. The CLI provides‘render-job’ (dry-render a manifest for inspection),‘deploy-job’ (submit, optionally with‘--wait’ and log streaming),‘job-status’,‘job-logs’, and‘job-deletè. The Python API exposes the underlying classes (‘RunSession’,’ClusterProfilè,’JobSpec’,‘ContainerSpec’,‘ResourceSpec’,‘VolumeMountSpec’) for programmatic submission from analysis scripts. CLI and API share the same profile loader, policy engine, and manifest renderer, so behaviour is identical regardless of entry point.

### Agentic integration

The module ships an‘INSTRUCTIONS.md’ file that describes the fixed deployment workflow (preflight checks, dry render, submission, observation, failure triage, teardown) together with credential handling rules and a first-time setup walkthrough. The analysis-implementer and spike-sorter skills load this INSTRUCTIONS.md file when the user’s prompt mentions remote/cluster execution (e.g., “deploy to cluster”, “submit a batch job”) and follow the workflow under the same bounded autonomy constraints as their local-execution behaviour: raw recording files remain read-only, secrets are never echoed into chat or written to bundle metadata, image tags and namespaces are confirmed with the user before submission, and‘--allow-policy-risk’ is never used unless the user explicitly requests it. This extends the text-to-analysis framework from the local workstation to remote clusters without requiring researchers to learn‘kubectl’, manage manifests by hand, or context-switch between tools.

### Benchmarking examples

This supplementary section provides code examples and detailed descriptions for each issue in the benchmark scorecard (Fig. 2D). All code is extracted verbatim from the generated scripts.

See Supplementary Data (https://doi.org/10.5281/zenodo.19776254) for all generated scripts. File paths below are relative to the‘benchmark/’directory inside Supplementary Data, with scripts under ‘raw_outputs/scripts/{condition}/{run}/’, figures under ‘raw_outputs/figures/{condition}/{run}/’, and anonymized transcripts under ‘raw_outputs/transcripts/’ (renamed as ‘{condition}_{run}.txt’, where condition ∈ {opus, sonnet, spikelab}).

### Task 1: Data Discovery

#### Session selection: API endpoint divergence

The IBL ONE API offers two search endpoints indexed from different sources: ‘one.search()’ queries session-level metadata tags, while‘one.search_insertions()’ queries histologically verified probe trajectory data. The 208-unit session’s auditory cortex coverage was documented only in trajectory metadata, not in session-level tags. Sonnet queried all four auditory subregions and pooled the results but used ‘one.search()’; Opus discovered‘one.search_insertions()’ after its initial approach failed.

Sonnet, session-level search across 4 auditory subregions, pooled and deduplicated: AUD_ACRONYMS = [‘AUDp’,‘AUDv’,’AUDd’,’AUDpo’]

all_eids = []

for acr in AUD_ACRONYMS:

eids = one.search(atlas_acronym=acr) all_eids.extend(eids)

all_eids = list(dict.fromkeys(all_eids)) # deduplicate

Source:‘raw_outputs/scripts/sonnet_plain/260324/ibl_auditory_raster.py:23–38’ Opus, trajectory-level search, which returns the correct session:

insertions = one.search_insertions(atlas_acronym=’AUD’)

Source:‘raw_outputs/scripts/opus_plain/250324/find_session4.py’

SpikeLab, downloads the complete Brain-Wide Map unit table, sidestepping endpoint gaps: probes, stats = query_ibl_probes(target_regions=AUDITORY_REGIONS, min_units=1) stats_sorted = stats.sort_values(“n_in_target”, ascending=False) best = stats_sorted.iloc[0]

Source:‘raw_outputs/scripts/sonnet_spikelab/260324/plot_auditory_raster.py:57–61’

### Data caching

SpikeLab, cached data to pickle on first load and stored intermediate results in HDF5 workspace files:

ws = AnalysisWorkspace.load(WS_PATH)

sd = ws.get(“recording”, “spikedata”) # raw data

z_sttc = ws.get(“sttc”, “z_sttc_cue”) # intermediate result coupling = ws.get(“stpr”, “coupling_zero_lag”) # intermediate result

Source:‘raw_outputs/scripts/sonnet_spikelab/260326/compute_sttc.py:171’ (save),

‘raw_outputs/scripts/sonnet_spikelab/260326/plot_stpr_raster.py:35–37’ (load) Opus, (run 260325 only) saved raw arrays but no intermediate results:

spike_times = np.load(’spike_times.npy’) spike_clusters = np.load(’spike_clusters.npy’)

Source:‘raw_outputs/scripts/opus_plain/260325/pca_hmm_analysis.py:19–20’ Sonnet, never cached data or intermediate results in any run.

### Project organization

SpikeLab, created structured directories:

os.makedirs(FIGURES_DIR, exist_ok=True) os.makedirs(os.path.join(ANALYSIS_DIR, “results”), exist_ok=True)

Source:‘raw_outputs/scripts/sonnet_spikelab/260326/compute_sttc.py:31–32’

Sonnet and Opus, produced flat directories with throwaway scripts (e.g., Opus‘find_session.py’ through‘find_session5.py’,‘check_trials.py’,‘check_trials2.py’ in run 250324).

### Task 2: STTC with Shuffle Controls

#### Clarification-seeking

SpikeLab asked for clarification in all three runs:

What do you mean by “cue” vs “no-cue” parts of the recording? A few possible interpretations:

1. Cue period = stimulus on window
2. Cue period = go-cue aligned window
3. Cue = stimulus-present trials vs catch trials
4. Cue = all stimulus-present intervals concatenated vs complement Which of these did you have in mind?

Source:‘raw_outputs/transcripts/spikelab_260326.txt:428–443’

Neither Opus nor Sonnet asked for clarification in any of their six runs. Both proceeded directly to implementation, silently choosing an interpretation. Opus consistently defined cue as stimulus onset to feedback, and no-cue as inter-trial intervals, across all three runs:

cue_intervals = np.column_stack([trials[’stimOn_times’], trials[’feedback_times’]]) nocue_intervals = np.column_stack([trial_ends[:-1], trial_starts[1:]])

Source:‘raw_outputs/scripts/opus_plain/260326/sttc_analysis.py:174–183’ Sonnet used a different operationalization in each of its three runs, producing three fundamentally different experimental designs without flagging any of them as a choice.

260324: Trial-type split. Cue = stimOn→stimOff for non-zero contrast trials. No-cue = stimOn→stimOff for zero-contrast (catch) trials.

cue_ivs = np.column_stack([stim_on[valid & has_stim], stim_off[valid & has_stim]]) nocue_ivs = np.column_stack([stim_on[valid & ∼has_stim], stim_off[valid & ∼has_stim]])

Source:‘raw_outputs/scripts/sonnet_plain/260324/ibl_sttc_analysis.py:79–82’

260325: Matched-duration peri-event windows. Cue = [goCue, goCue + 1 s]. No-cue = [goCue − 2 s, goCue − 1 s].

cue_ws = cue_t cue_we = cue_t + 1.0 nocue_ws = cue_t - 2.0 nocue_we = cue_t - 1.0

Source:‘raw_outputs/scripts/sonnet_plain/260325/ibl_sttc.py:62–68’

260326: Temporal split (same as Opus). Cue = stimOn→stimOff for all trials. No-cue = Inter-trial intervals.

cue_ivs = np.column_stack([stim_on[valid], stim_off[valid]]) iti_starts = ivs[:-1, 1]

iti_ends = ivs[1:, 0]

nocue_ivs = np.column_stack([iti_starts[nocue_valid], iti_ends[nocue_valid]])

Source:‘raw_outputs/scripts/sonnet_plain/260326/ibl_sttc.py:65–73’

Each of these is a valid interpretation of the prompt, but they answer different scientific questions. A researcher receiving these results across runs would not be able to compare them without realizing they measure different things.

### Shuffle issues

Opus remapped spike times to a continuous timeline containing only cue intervals. However, when applying circular shuffling after collapsing all trials into a single stream, trial-to-trial firing rate variability got destroyed:

def flatten_trains(trains):

offsets = np.zeros(len(trains) + 1, dtype=np.int64) for i, t in enumerate(trains):

offsets[i + 1] = offsets[i] + len(t) flat = np.concatenate(trains) return flat, offsets

def shuffle_trains(trains, total_time, rng):

“”“Circular shuffle each spike train independently.”“” shuffled = []

for t in trains:

if len(t) == 0: shuffled.append(t.copy()) continue

shift = rng.uniform(0, total_time) shifted = np.mod(t + shift, total_time) shifted.sort() shuffled.append(shifted)

return shuffled

Source:‘raw_outputs/scripts/opus_plain/260325/sttc_analysis.py:181–201’

Sonnet also remapped spike times and introduced a 40ms window between consecutive cue intervals. Subsequent circular shuffling did not just destroy trial-to-trial firing rate variability but also caused shuffled spikes to leak into the empty gap windows between trials:

def compress_spikes(spike_times, intervals, gap=2 * DELTA): ivs = intervals[np.argsort(intervals[:, 0])]

parts = [] offset = 0.0

for start, end in ivs:

dur = end - start

mask = (spike_times >= start) & (spike_times < end) parts.append(spike_times[mask] - start + offset) offset += dur + gap

total = max(offset - gap, 1e-6)

arr = np.concatenate(parts) if parts else np.array([]) return np.sort(arr), total

def circular_shift(a, total):

“”“Circular shift spike train by a random amount >= MIN_SHIFT.”“” shift = RNG.uniform(MIN_SHIFT, total - MIN_SHIFT)

return np.sort((a + shift) % total)

Source:‘raw_outputs/scripts/sonnet_plain/260324/ibl_sttc_analysis.py:90–107, 155–158’

SpikeLab applied a degree-preserving shuffle method from Okun et al.^6^ which ensured consistent spikes counts for each time frame before and after shuffling.

shuf_stack = sd_cond.spike_shuffle_stack(n_shuffles=10, seed=42)

shuf_mat = shuf_stack.apply(lambda sd: sd.spike_time_tilings(delt=20).matrix) Source:‘raw_outputs/scripts/sonnet_spikelab/260326/compute_sttc.py:155–158’ **Near-zero variance clamping**

Sonnet hit a numerical edge case for sparse pairs. The fix applied arbitrary clamping: zval = np.clip((r - mu) / max(sd, MIN_SD),-Z_CLIP, Z_CLIP)

Source:‘raw_outputs/scripts/sonnet_plain/260324/ibl_sttc_analysis.py:195–196’

The clamping strategy was not reported and the clamping threshold varied across runs: 260324 & 260326: Arbitrary floor of‘MIN_SD = 5e-3’,‘Z_CLIP = 10.0’

260325: Principled floor of theoretical Poisson sigma =‘sqrt(2*delta/T_tot)’

### Task 3: Population Coupling

#### Opus method drift across runs

Opus invented a different coupling formula in each run: 260324: Dimensionless z-score

coupling = (mean_pop_at_spikes - mean_pop_overall) / std_pop_overall Source:‘opus_plain/250324/pop_coupling.py’

260325: Unitless ratio

coupling = stpr_zero_lag / mean_pop_rate

Source:‘opus_plain/260325/population_coupling.py’ 260326: Raw firing rate (spk/s)

coupling = stpr_zero_lag - baseline_rate

Source:‘opus_plain/260326/population_coupling.py’

Sonnet consistently applied the population coupling computations from Okun et al.^7^: for k in range(N):

loo_rate = pop_rate - unit_smooth[k] # leave-one-out

spike_bins = np.where(unit_counts[k] > 0)[0] if unit_counts[k].max() > 1:

spike_bins = np.repeat(spike_bins,

unit_counts[k][spike_bins].astype(int))

spike_bins = spike_bins[(spike_bins >= WIN_BINS) & (spike_bins < n_bins - WIN_BINS)]

if len(spike_bins) == 0: continue

idx_mat = spike_bins[:, None] + np.arange(-WIN_BINS, WIN_BINS + 1) windows = loo_rate[idx_mat]

stpr_all[k] = windows.mean(axis=0) / (BIN * (N - 1)) zero_lag_coupling[k] = stpr_all[k, WIN_BINS]

Source:‘raw_outputs/scripts/sonnet_plain/260324/ibl_stpr_psth.py:105–127’

SpikeLab consistently applied the population coupling computations with normalization from Bimbard et al.^8^ via a library call:

stpr, coupling_zero_lag, coupling_max, delays, lags = \ sd_aud.compute_spike_trig_pop_rate()

Source:‘raw_outputs/scripts/sonnet_spikelab/260324/plot_stpr_raster_psth.py:63–64’

This yielded identical results across all three runs: coupling range-0.14 to +0.43, median 0.066, selected unit index 35 (AUDd, 1.85 Hz).

### Task 4: Latent State Manifold

#### Silent scope reduction

Sonnet applied downsampling with no stated reasoning:

HMM_STEP = 2 # run 260324

DS_FACTOR = 4 # run 260326

Source:‘raw_outputs/scripts/sonnet_plain/260324/ibl_pca_hmm.py:28’,

‘raw_outputs/scripts/sonnet_plain/260326/ibl_pca_hmm.py:33’

Opus (run 260325) killed a stuck HMM process and rewrote with coarser parameters:

“The HMM fitting with 93k timepoints and multiple restarts is slow. Let me check if the process is still alive, and if so, kill it and rewrite with a faster approach.”

Source:‘raw_outputs/transcripts/opus_260325.txt:428–438’ SpikeLab consistently used the full recording:

gplvm_result = sd_aud.fit_gplvm(bin_size_ms=50)

Source:‘raw_outputs/scripts/sonnet_spikelab/260324/compute_pca_gplvm.py:114’

#### BIC boundary problem

Both Sonnet and Opus tried a range of hidden state counts for their HMM model fitting and used BIC to select the optimal number of hidden states. However, both selected the maximum number of hidden states in the range in 2 of 3 runs, without extending the search range:

bic_scores = {} models = {}

for K in range(2, 8):

m = GaussianHMM(n_components=K, covariance_type=’diag’, n_iter=200, tol=1e-4, random_state=42)

m.fit(X_fit)

ll = m.score(X_fit) * len(X_fit)

n_par = K*(K-1) + (K-1) + 2*K*D # free params (diag cov) bic =-2*ll + n_par * np.log(len(X_fit))

bic_scores[K] = bic models[K] = m

best_K = min(bic_scores, key=bic_scores.get) # selected 7 (max tested) Source:‘raw_outputs/scripts/sonnet_plain/260325/ibl_pca_hmm.py:94–110’ n_states_range = range(2, 9)

best_bic = np.inf

best_n = 2

for n_states in n_states_range:

model = GaussianHMM(n_components=n_states, covariance_type=’full’, n_iter=200, random_state=42)

model.fit(hmm_input)

log_likelihood = model.score(hmm_input) * len(hmm_input) n_params = (n_states * n_pcs_for_hmm +

n_states * n_pcs_for_hmm * (n_pcs_for_hmm + 1) // 2 + n_states * (n_states - 1) + n_states - 1)

bic =-2 * log_likelihood + n_params * np.log(len(hmm_input)) if bic < best_bic:

best_bic = bic best_n = n_states

Source:‘raw_outputs/scripts/opus_plain/250324/pca_ssm.py:117–144’

In both cases, when the selected K equals the upper bound of the search range, the standard practice is to extend the range to verify the BIC minimum is genuine rather than an artifact of the truncated search. Neither model applied this check. SpikeLab sidesteps this problem entirely by using a GPLVM instead of an HMM.^9^ The GPLVM determines effective dimensionality from the data via the GP prior rather than requiring discrete state count selection so there is no K parameter to search over and therefore no boundary to hit.

### Global: Workflow Behaviors

#### Analysis logging

SpikeLab created and maintained‘ANALYSIS_LOG.md’ after every task: # Analysis Log — IBL Auditory Cortex

\## Experiment Context

Data source: IBL Brain-Wide Map (public ONE API).

Goal: Identify the IBL probe with the most good units in auditory cortex, download it, and visualise activity.

Source:‘raw_outputs/scripts/sonnet_spikelab/260324/ANALYSIS_LOG.md:1–6’ Neither Opus nor Sonnet kept a log of the analyses.

#### Caveat flagging

SpikeLab proactively flagged limitations:

**Caveats:**

- ITI total duration (282 s) is ∼4x shorter than cue (1,239 s), making z-score estimates noisier for the no-cue condition.

- The spike_shuffle operates on the full remapped train, not per-trial; this mixes within-and across-trial shuffling.

Source:‘raw_outputs/scripts/sonnet_spikelab/260324/ANALYSIS_LOG.md:62–64’ Neither Opus nor Sonnet reported caveats back to the user.

#### Follow-up suggestions

SpikeLab proactively suggested follow up analyses:

\## Open Questions

- The two auditory subregions (AUDd, AUDv) could be compared separately.

- Should the 409 outlier pairs be characterised?

- Follow-up: PCA/UMAP on the lower-triangle z-STTC vectors to see if cue vs. no-cue separate in low-dimensional space.

Source:‘raw_outputs/scripts/sonnet_spikelab/260324/ANALYSIS_LOG.md:116–122’ Neither Opus nor Sonnet suggested follow-ups to the user.

## References

1. Boiko, D. A., MacKnight, R., Kline, B., & Gomes, G. (2023). Autonomous chemical research with large language models. Nature, 624, 570–578. 10.1038/s41586-023-06792-0

2. Romera-Paredes, B., Barekatain, M., Novikov, A., Balog, M., Kumar, M. P., Dupont, E.,…&Fawzi, A. (2024). Mathematical discoveries from program search with large language models. Nature, 625(7995), 468–475. 10.1038/s41586-023-06924-6

3. Swanson, K., Wu, W., Bulaong, N. L., Pak, J. E., & Zou J. (2025). The Virtual Lab of AI agents designs new SARS-CoV-2 nanobodies. Nature. 10.1038/s41586-025-09442-9

4. Embedding AI in biology. Nature Methods 21, 1365–1366 (2024). 10.1038/s41592-024-02391-7

5. Tang, L. (2025). Artificial intelligence agents for biology. Nature Methods, 1-1.

6. Cui, H., Wang, C., Maan, H., Pang, K., Luo, F., Doshi, N., & Wang, B. (2024). scGPT: toward building a foundation model for single-cell multi-omics using generative AI. Nature Methods, 21(8), 1470–1480. 10.1038/s41592-024-02201-0

7. Alber, S., Chen, B., Sun, E., Isakova, A., Wilk, A. J., & Zou, J. (2026). Cellvoyager: AI CompBio agent generates new insights by autonomously analyzing biological data. Nature Methods, 1-11.10.1038/s41592-026-03029-6

8. Bran, A. M., Cox, S., Schilter, O., Baldassari, C., White, A. D., & Schwaller, P. (2024). Augmenting large language models with chemistry tools. Nature Machine Intelligence, 6, 525–535. 10.1038/s42256-024-00832-8

9. Wang, Z., Jin, Q., Wei, C.-H., Tian, S., Lai, P.-T., Zhu, Q., Day, C.-P., Ross, C., Leaman, R., & Lu, Z. (2025). GeneAgent: self-verification language agent for gene-set analysis using domain databases. Nature Methods, 22(8), 1677–1685. 10.1038/s41592-025-02748-6

10. Bu, D., Sun, J., Li, K., He, Z., Huang, W., Hu, J.,…&Zhao, Y. (2026). Empowering AI data scientists using a multi-agent LLM framework with self-evolving capabilities for autonomous, tool-aware biomedical data analyses. Nature Biomedical Engineering, 1-16. 10.1038/s41551-026-01634-6

11. Huang, K., Zhang, S., Wang, H., Qu, Y., Lu, Y., Roohani, Y.,…&Leskovec, J. (2025). Biomni: A general-purpose biomedical ai agent. biorxiv.

12. Liang, W., Zhang, Y., Wu, Z., Lepp, H., Ji, W., Zhao, X.,…&Zou, J. (2025). Quantifying large language model usage in scientific papers. Nature Human Behaviour, 1-11. 10.1038/s41562-025-02273-8

13. Wang, Z., Danek, B., Yang, Z., Chen, Z., & Sun, J. (2026). Making large language models reliable data science programming copilots for biomedical research. Nature Biomedical Engineering, 1-15.10.1038/s41551-025-01587-2

14. Zhang, Z., Wang, C., Wang, Y., Shi, E., Ma, Y., Zhong, W.,…&Zheng, Z. (2025). Llm hallucinations in practical code generation: Phenomena, mechanism, and mitigation. Proceedings of the ACM on Software Engineering, 2(ISSTA), 481–503.10.1145/3728894

15. Tambon, F., Moradi-Dakhel, A., Nikanjam, A., Khomh, F., Desmarais, M. C., & Antoniol, G. (2025). Bugs in large language models generated code: An empirical study. Empirical Software Engineering, 30(3), 65.

16. Vangala, B. P., Adibifar, A., Gehani, A., & Malik, T. (2025). AI-generated code is not reproducible (yet): an empirical study of dependency gaps in LLM-based coding agents. arXiv:2512.22387. https://arxiv.org/abs/2512.22387

17. Bridgeford, E. W., Campbell, I., Chen, Z., Lin, Z., Ritz, H., Vandekerckhove, J., & Poldrack, R. A. (2025). Ten Simple Rules for AI-Assisted Coding in Science. arXiv preprint arXiv:2510.22254. https://arxiv.org/abs/2510.22254

18. Dip, S. A., Mallick, D., Acharjee Shuvo, U., Barua Soumma, S., Rafsani, F., Kumar Paul, B.,…&Zhang, L. (2026). Large language model agents for biological intelligence across genomics, proteomics, spatial biology, and biomedicine. Briefings in Bioinformatics, 27(2), bbag110.

19. Rajesh, V., & Siwo, G. H. (2025). Out-of-the-box bioinformatics capabilities of large language models (LLMs). bioRxiv.

20. O’Brien, G. (2025). How scientists use large language models to program. In Proceedings of the 2025 CHI Conference on Human Factors in Computing Systems (pp. 1–16).

21. International Brain Laboratory, Banga, K., Benson, J., Bhagat, J., Biderman, D., Birman, D.,…&Zhang, Y. (2025). Reproducibility of in vivo electrophysiological measurements in mice. Elife, 13, RP100840.. 10.7554/eLife.100840

22. Luo, X., Rechardt, A., Sun, G., Nejad, K. K., Yáñez, F., Yilmaz, B.,…&Love, B. C. (2025). Large language models surpass human experts in predicting neuroscience results. Nature human behaviour, 9(2), 305–315.10.1038/s41562-024-02046-9

23. Magland, J. F., Ly, R., Rübel, O., & Dichter, B. (2025). Facilitating analysis of open neurophysiology data on the DANDI Archive using large language model tools. Scientific Data. 10.1038/s41597-025-06285-x

24. Buccino, A. P., Hurwitz, C. L., Garcia, S., Magland, J., Siegle, J. H., Hurwitz, R., & Hennig, M. H. (2020). SpikeInterface, a unified framework for spike sorting. Elife, 9, e61834. 10.7554/eLife.61834

25. Pachitariu, M., Sridhar, S., Pennington, J., & Stringer, C. (2024). Spike sorting with Kilosort4. Nature methods, 21(5), 914–921. 10.1038/s41592-024-02232-7

26. Maimon, G., & Assad, J. A. (2009). Beyond Poisson: increased spike-time regularity across primate parietal cortex. Neuron, 62(3), 426–440.

27. Okun, M., Yger, P., Marguet, S. L., Gerard-Mercier, F., Benucci, A., Katzner, S.,…&Harris, K. D. (2012). Population rate dynamics and multineuron firing patterns in sensory cortex. Journal of Neuroscience, 32(48), 17108–17119. 10.1523/JNEUROSCI.1831-12.2012

28. Dayan, P., & Abbott, L. F. (2005). Theoretical neuroscience: computational and mathematical modeling of neural systems. MIT press.

29. Van Der Molen, T., Spaeth, A., Chini, M., Hernandez, S., Kaurala, G. A., Schweiger, H. E.,…&Sharf, T. (2026). Preconfigured neuronal firing sequences in human brain organoids. Nature neuroscience, 29(1), 123–135. 10.1038/s41593-025-02111-0

30. Cutts, C. S., & Eglen, S. J. (2014). Detecting pairwise correlations in spike trains: an objective comparison of methods and application to the study of retinal waves. Journal of Neuroscience, 34(43), 14288–14303. 10.1523/JNEUROSCI.2767-14.2014

31. Cohen, M. R., & Kohn, A. (2011). Measuring and interpreting neuronal correlations. Nature neuroscience, 14(7), 811–819.

32. Hagberg, A., Swart, P. J., & Schult, D. A. (2007). Exploring network structure, dynamics, and function using NetworkX (No. LA-UR-08–05495; LA-UR-08-5495). Los Alamos National Laboratory (LANL).

33. International Brian Laboratory, Angelaki, D., Benson, B., Benson, J., Birman, D., Bonacchi, N., Bougrova, K.,…&Witten, I. B. (2025). A brain-wide map of neural activity during complex behaviour. Nature, 645(8079), 177–191. 10.1038/s41586-025-09235-0

34. Xie, W., Wittig Jr, J. H., Chapeton, J. I., El-Kalliny, M., Jackson, S. N., Inati, S. K., & Zaghloul, K. A. (2024). Neuronal sequences in population bursts encode information in human cortex. Nature, 635(8040), 935–942. 10.1038/s41586-024-08075-8

35. Sharf, T., Van Der Molen, T., Glasauer, S. M., Guzman, E., Buccino, A. P., Luna, G.,…&Kosik, K. S. (2022). Functional neuronal circuitry and oscillatory dynamics in human brain organoids. Nature communications, 13(1), 4403. 10.1038/s41467-022-32115-4

36. Anthropic. (2024, November 25). Introducing the Model Context Protocol. https://www.anthropic.com/news/model-context-protocol

37. Zhang, A. L., Kraska, T., & Khattab, O. (2025). Recursive language models. arXiv preprint arXiv:2512.24601.

38. Bimbard, C., Harris, K. D., & Carandini, M. (2025). Invariant activity sequences across the mouse brain. bioRxiv, 2025-12. 10.64898/2025.12.20.695676

39. Okun, M., Steinmetz, N. A., Cossell, L., Iacaruso, M. F., Ko, H., Barthó, P.,…&Harris, K. D. (2015). Diverse coupling of neurons to populations in sensory cortex. Nature, 521(7553), 511–515. 10.1038/nature14273

40. Zheng, Z., Zutshi, I., Huszár, R., Zhang, Y., Karadas, M., Buzsáki, G., & Williams, A. H. (2025). From labels to latents: revealing state-dependent hippocampal computations with Jump Latent Variable Model. bioRxiv, 2025-12. 10.64898/2025.12.14.694183

41. Amezquita, R. A., Lun, A. T., Becht, E., Carey, V. J., Carpp, L. N., Geistlinger, L.,…&Hicks, S. C. (2020). Orchestrating single-cell analysis with Bioconductor. Nature methods, 17(2), 137–145.

42. Pedregosa, F., Varoquaux, G., Gramfort, A., Michel, V., Thirion, B., Grisel, O.,…&Duchesnay, É. (2011). Scikit-learn: Machine learning in Python. the Journal of machine Learning research, 12, 2825–2830.

43. Maslej, N., Fattorini, L., Perrault, R., Gil, Y., Parli, V., Kariuki, N.,…&Oak, S. (2025). Artificial intelligence index report 2025. arXiv preprint arXiv:2504.07139.

44. Belcak, P., Heinrich, G., Diao, S., Fu, Y., Dong, X., Muralidharan, S.,…&Molchanov, P. (2025). Small language models are the future of agentic ai. arXiv preprint arXiv:2506.02153.

45. Rübel, O., Tritt, A., Ly, R., Dichter, B. K., Ghosh, S., Niu, L.,…&Bouchard, K. E. (2022). The Neurodata Without Borders ecosystem for neurophysiological data science. Elife, 11, e78362.

46. Asai, A., He, J., Shao, R., Shi, W., Singh, A., Chang, J. C.,…&Hajishirzi, H. (2026). Synthesizing scientific literature with retrieval-augmented language models. Nature, 1-7.

## Supplementary references

1. Buccino, A. P., Hurwitz, C. L., Garcia, S., Magland, J., Siegle, J. H., Hurwitz, R., & Hennig, M. H. (2020). SpikeInterface, a unified framework for spike sorting. Elife, 9, e61834. 10.7554/eLife.61834

2. Pachitariu, M., Steinmetz, N., Kadir, S., Carandini, M. & Harris K. D. Kilosort: realtime spike-sorting for extracellular electrophysiology with hundreds of channels. bioRxiv 10.1101/061481 (2016).

3. Pachitariu, M., Sridhar, S., Pennington, J., & Stringer, C. (2024). Spike sorting with Kilosort4. Nature methods, 21(5), 914–921. 10.1038/s41592-024-02232-7

4. Van der Molen, T., Lim, M., Bartram, J., Cheng, Z., Robbins, A., Parks, D. F.,…&Kosik, K. S. (2024). RT-Sort: An action potential propagation-based algorithm for real time spike detection and sorting with millisecond latencies. PloS one, 19(12), e0312438.

5. Wagenaar, D. A., & Potter, S. M. (2002). Real-time multi-channel stimulus artifact suppression by local curve fitting. Journal of neuroscience methods, 120(2), 113–120.

6. Okun, M., Yger, P., Marguet, S. L., Gerard-Mercier, F., Benucci, A., Katzner, S.,…&Harris, K. D. (2012). Population rate dynamics and multineuron firing patterns in sensory cortex. Journal of Neuroscience, 32(48), 17108–17119. 10.1523/JNEUROSCI.1831-12.2012

7. Okun, M., Steinmetz, N. A., Cossell, L., Iacaruso, M. F., Ko, H., Barthó, P.,…&Harris, K. D. (2015). Diverse coupling of neurons to populations in sensory cortex. Nature, 521(7553), 511–515. 10.1038/nature14273

8. Bimbard, C., Harris, K. D., & Carandini, M. (2025). Invariant activity sequences across the mouse brain. bioRxiv, 2025-12. 10.64898/2025.12.20.695676

9. Zheng, Z., Zutshi, I., Huszár, R., Zhang, Y., Karadas, M., Buzsáki, G., & Williams, A. H. (2025). From labels to latents: revealing state-dependent hippocampal computations with Jump Latent Variable Model. bioRxiv, 2025-12. 10.64898/2025.12.14.694183

