## Supplementary material for "SpikeLab: Agentic tools for spike data analysis": Spikelab skill: INSTALL.md

# SpikeLab Installation Guide

There are two installation paths:

- **PyPI install** — for users who only want to *use* the library and the agent skills (analysis, education, spike sorting). No source clone required.
- **Source clone + editable install** — required for the developer skill, for contributing PRs, or any workflow that edits library source code.

The agent skills (analysis-implementer, educator, map-updater, developer, spikesorter) ship as package data, so they are present after either install path.

---

## Quick install from PyPI

```bash
pip install spikelab
```

Extras work the same way as the editable install below:

```bash
pip install "spikelab[s3]"
pip install "spikelab[s3,ml,mcp]"   # multiple extras
pip install "spikelab[all]"         # everything except kilosort4
```

The full extras list is identical to the editable install — see the table further down.

After install, the agent skills live at:

```bash
python -c "import spikelab; print(spikelab.__path__[0])"
# → <env>/site-packages/spikelab/
# Skills are under: <env>/site-packages/spikelab/agent/skills/
```

The map-updater writes `REPO_MAP.md` and `REPO_MAP_DETAILED.md` next to its own SKILL.md inside `site-packages/`. They will be wiped by `pip install --upgrade spikelab` and need to be regenerated. This is by design.

---

## Source clone + editable install

For development, contributing PRs, or any workflow that edits library source.

### Clone the repository

```bash
git clone https://github.com/braingeneers/SpikeLab.git
```

### Install the library

**Option A — Conda environment (recommended for full functionality):**

The repository includes an `environment.yml` that installs Python, all core dependencies, and all optional packages (matplotlib, scikit-learn, umap-learn, networkx, neo, boto3, etc.):

```bash
conda env create -f SpikeLab/environment.yml
conda activate spikelab
pip install -e SpikeLab/
```

This is the recommended approach because `umap-learn` depends on `numba`/`llvmlite`, which install cleanly from conda-forge but often fail when built from source via pip.

**Option B — Pip only (core + selected extras):**

Install the core library (numpy, scipy, matplotlib, h5py):

```bash
pip install -e SpikeLab/
```

Install with optional dependency groups using extras:

```bash
# Core + MCP server (mcp)
pip install -e "SpikeLab/[mcp]"

# Core + MCP SSE transport (uvicorn, starlette)
pip install -e "SpikeLab/[sse]"

# Core + S3 support (boto3)
pip install -e "SpikeLab/[s3]"

# Core + extra I/O helpers (pandas)
pip install -e "SpikeLab/[io]"

# Core + ML (scikit-learn, umap-learn, networkx, python-louvain)
pip install -e "SpikeLab/[ml]"

# Core + Neo/NWB interop (neo, quantities, pynwb)
pip install -e "SpikeLab/[neo]"

# Core + GPLVM latent state modelling (jax, jaxlib, jaxopt, optax, poor-man-gplvm)
pip install -e "SpikeLab/[gplvm]"

# Core + numba-accelerated kernels
pip install -e "SpikeLab/[numba]"

# Core + Kilosort2 / rt-sort spike sorting (spikeinterface, natsort, six, pandas)
pip install -e "SpikeLab/[spike-sorting]"

# Core + Kilosort4 (kilosort — also requires PyTorch with CUDA, installed separately)
pip install -e "SpikeLab/[kilosort4]"

# Core + Kubernetes batch jobs (pydantic, PyYAML, Jinja2, kubernetes)
pip install -e "SpikeLab/[batch-jobs]"

# Core + docs build (sphinx, sphinx-rtd-theme, sphinx-autodoc-typehints)
pip install -e "SpikeLab/[docs]"

# Core + dev tools (pytest, pytest-asyncio, black)
pip install -e "SpikeLab/[dev]"

# Everything (all of the above except kilosort4)
pip install -e "SpikeLab/[all]"
```

Multiple extras can be combined: `pip install -e "SpikeLab/[s3,ml,mcp]"`.

Note: `matplotlib` is a core dependency, so plotting works without any extra. There is no `[plotting]` extra.

Note: `umap-learn` depends on `numba`/`llvmlite`. If pip installation fails, install via conda instead: `conda install -c conda-forge umap-learn`.

### Updating SpikeLab (source clone)

```bash
cd SpikeLab && git pull && cd ..
pip install -e SpikeLab/
```

Or if using the conda environment: `conda env update -f SpikeLab/environment.yml` followed by `pip install -e SpikeLab/`.

After pulling new code, regenerate the repo maps (see below).

For PyPI installs, update with `pip install --upgrade spikelab` and then regenerate the repo maps (the upgrade wipes them).

---

## Generating the Repo Maps (required)

The repo maps do not ship with the package — they must be generated after installation and after every library update. Without them, the analysis-implementer, educator, developer, and spike-sorter skills cannot orient themselves to the library API and will produce unreliable results.

Run the `spikelab-map-updater` skill to generate `REPO_MAP.md` and `REPO_MAP_DETAILED.md`. The skill ships inside the spikelab package at:

```
agent/skills/spikelab-map-updater/SKILL.md
```

To find the package directory (works for both editable and PyPI installs):

```bash
python -c "import spikelab; print(spikelab.__path__[0])"
# editable: <clone>/SpikeLab/src/spikelab
# PyPI:     <env>/site-packages/spikelab
```

The skill reads the library source from there and produces two files in the same `agent/skills/spikelab-map-updater/` directory:
- `REPO_MAP.md` — condensed quick reference
- `REPO_MAP_DETAILED.md` — full API reference with signatures and return types

**When to regenerate:**
- After first installation
- After `git pull` with library changes (editable install)
- After `pip install --upgrade spikelab` (PyPI install — the upgrade wipes the maps)
- After adding or modifying methods in the library
- If any skill reports that the repo maps are missing or outdated

**Note for PyPI installs:** the maps are written into `site-packages/`, which requires that the environment is user-writable. This is fine for venv and conda environments; system Python installs would need `--user` or sudo.
