## Supplementary material for "SpikeLab: Agentic tools for spike data analysis": Spikelab skill: SKILL.md

---
name: spikelab
description: Sets up and manages the SpikeLab environment. Clones the repository, installs the library, generates repo maps, and delegates to in-repo skills for analysis, spike sorting, development, and education. Use when the user wants to work with SpikeLab.
---

# SpikeLab

SpikeLab is a Python library for loading, analyzing, visualizing, and exporting neuronal spike train data from multi-electrode array (MEA) electrophysiology experiments.

---

## Session Start — Environment Detection

Run this detection sequence once at the start of every session:

1. Check if `spikelab` is importable in the default `python`:
   ```bash
   python -c "import spikelab; print('OK')" 2>/dev/null
   ```
   If this succeeds, use bare `python` for all commands.

2. If it fails, check for a `spikelab` conda environment:
   ```bash
   conda env list 2>/dev/null | grep spikelab
   ```
   If found, prefix all commands with `conda run -n spikelab` for the rest of the session.

3. If neither works, **read `INSTALL.md`** in this skill directory for full installation instructions. Prefer the conda approach (Option A) unless the user specifies otherwise.

Use the detected invocation method consistently for all Python/pip commands in the session.

---

## Delegation

Once SpikeLab is available, delegate to the appropriate in-repo skill based on the user's request:

**Spike sorting** — the user wants to sort raw recordings, configure sorting parameters, curate units, assess sorting quality, or troubleshoot sorting failures:
```
SpikeLab/src/spikelab/agent/skills/spikelab-spikesorter/SKILL.md
```
It covers: `sort_recording`, `sort_multistream`, curation, QC figures, and loading/inspecting sorted results. Outputs are saved as `sorted_spikedata_curated.pkl` files that the analysis implementer can load.

**Explanatory and conceptual questions** — the user asks what an analysis does, how a method works, what a result means, wants to understand the underlying neuroscience, or asks about SpikeLab's API and data structures:
```
SpikeLab/src/spikelab/agent/skills/spikelab-educator/SKILL.md
```
It contains concept definitions, interpretation guidance, literature references, and cross-references to documentation, examples, and source code.

**Library development** — the user wants to integrate analysis code into the library, promote a script's computations to reusable methods, add new functionality, write tests, or expose methods through MCP:
```
SpikeLab/src/spikelab/agent/skills/spikelab-developer/SKILL.md
```
It covers the full integration pipeline: auditing scripts against existing methods, replacing reimplementations, adding novel methods, writing tests (main usage + edge cases), MCP exposure, and PR submission.

**Data analysis and visualization** — loading sorted data, running analyses, producing figures, writing scripts, and any ambiguous requests that are not sorting-related or development-related:
```
SpikeLab/src/spikelab/agent/skills/spikelab-analysis-implementer/SKILL.md
```
It contains the full analysis workflow: API orientation via repo maps, script structure, workspace usage, figure conventions, and reporting. It knows how to load sorting outputs from `sorted_spikedata_curated.pkl` files.

---

## Repo Maps

The repo maps do not ship with the repository — they must be generated after installation and after every library update. Without them, the analysis-implementer, educator, developer, and spike-sorter skills cannot orient themselves to the library API and will produce unreliable results.

Run the `spikelab-map-updater` skill to generate `REPO_MAP.md` and `REPO_MAP_DETAILED.md`:
```
SpikeLab/src/spikelab/agent/skills/spikelab-map-updater/SKILL.md
```

---

## Quick Reference

| What | Where |
|---|---|
| Library source | `SpikeLab/src/spikelab/` |
| Repo maps (API docs) | `SpikeLab/src/spikelab/agent/skills/spikelab-map-updater/REPO_MAP.md`, `REPO_MAP_DETAILED.md` |
| Map updater skill | `SpikeLab/src/spikelab/agent/skills/spikelab-map-updater/SKILL.md` |
| In-repo spike sorter skill | `SpikeLab/src/spikelab/agent/skills/spikelab-spikesorter/SKILL.md` |
| In-repo developer skill | `SpikeLab/src/spikelab/agent/skills/spikelab-developer/SKILL.md` |
| In-repo analysis skill | `SpikeLab/src/spikelab/agent/skills/spikelab-analysis-implementer/SKILL.md` |
| In-repo educator skill | `SpikeLab/src/spikelab/agent/skills/spikelab-educator/SKILL.md` |
| Installation guide | `INSTALL.md` (this skill directory) |
